# Reentrant DNA shells tune polyphosphate condensate size

**DOI:** 10.1101/2023.09.13.557044

**Authors:** Ravi Chawla, Jenna K. A. Tom, Tumara Boyd, Danielle A. Grotjahn, Donghyun Park, Ashok A. Deniz, Lisa R. Racki

## Abstract

The ancient, inorganic biopolymer polyphosphate (polyP) occurs in all three domains of life and affects myriad cellular processes. An intriguing feature of polyP is its frequent proximity to chromatin, and in the case of many bacteria, its occurrence in the form of magnesium-enriched condensates embedded in the nucleoid, particularly in response to stress. The physical basis of the interaction between polyP and DNA, two fundamental anionic biopolymers, and the resulting effects on the organization of both the nucleoid and polyP condensates remain poorly understood. Given the essential role of magnesium ions in the coordination of polymeric phosphate species, we hypothesized that a minimal system of polyP, magnesium ions, and DNA (polyP-Mg^2+^-DNA) would capture key features of the interplay between the condensates and bacterial chromatin. We find that DNA can profoundly affect polyP-Mg^2+^ coacervation even at concentrations several orders of magnitude lower than found in the cell. The DNA forms shells around polyP-Mg^2+^ condensates and these shells show reentrant behavior, primarily forming in the concentration range close to polyP-Mg^2+^ charge neutralization. This surface association tunes both condensate size and DNA morphology in a manner dependent on DNA properties, including length and concentration. Our work identifies three components that could form the basis of a central and tunable interaction hub that interfaces with cellular interactors. These studies will inform future efforts to understand the basis of polyP granule composition and consolidation, as well as the potential capacity of these mesoscale assemblies to remodel chromatin in response to diverse stressors at different length and time scales.

## INTRODUCTION

Polyphosphate (polyP) is a structurally simple, inorganic polymer consisting of a few to many hundreds of orthophosphate units linked by phosphoanhydride bonds. Biosynthesis of polyP is found in all three domains of life, and affects myriad cellular processes. In bacteria, polyP has been implicated in promoting cellular fitness with pleiotropic effects on biofilm formation, motility, cell cycle, and oxidative stress resistance^1–4^. In eukaryotic organisms, including humans, polyP has been linked with a wide variety of cellular processes from blood clotting and innate immunity to mitochondrial bioenergetics and cancer signaling^5,6^. How synthesis of this simple polyanion exerts a broad range of effects on cellular physiology has remained enigmatic. A major challenge to determining its molecular function has long been identifying and validating molecular interaction partners. While lacking known specificity epitopes at the primary level of organization, the polymer forms membraneless condensates in many bacteria that are spatially and temporally organized^7–11^.

A unifying organizational feature of polyP across evolution is that this polymer is frequently observed in close proximity with chromatin. In eukaryotes, from yeast to protists to metazoans, including human cells, polyP has been found in the nucleus, and also in some cases in the nucleolus^12–19^. Although the spatial organization of these granules differs within bacterial species, a longstanding and curious observation is that polyP granules associate with the nucleoid in many species. Embedding of polyphosphate granules within the nucleoid of diverse bacterial taxa has been observed at least since the 1960s^7,9,10,20–22^. In the opportunistic human pathogen *Pseudomonas aeruginosa*, polyP granules are transiently evenly spaced on the long axis of the cell in the nucleoid region^10^. In *Caulobacter crescentus*, polyP granules form at the ¼ and ¾ positions in the nucleoid region, and disruption of chromosome segregation can alter the granule organization, suggesting a functional association^7^.

In addition to this structural association, functional coupling between PolyP granules and DNA has been noted across different bacterial species. In *C. crescentus* and *E. coli* polyP synthesis affects cell cycle progression, and in *P. aeruginosa*, polyP promotes efficient cell cycle exit during starvation^10,23,24^. During nitrogen starvation, the SOS DNA damage response is activated in *P. aeruginosa* cells unable to make polyP, suggesting that polyP promotes nucleoid integrity by unknown mechanisms^10^. Recent work also demonstrates that polyphosphate drives heterochromatin formation in *E. coli* by modulating the DNA-binding affinity of nucleoid associated proteins (NAPs) like Hfq^25^. We previously found that polyP granules in *P. aeruginosa* are enriched in specific DNA binding proteins, including the histone H1-like protein AlgP^26^. Together, these observations implicate polyP condensates as an important feature of bacterial chromatin.

To address this hypothesis, that polyP condensates are a fundamentally important feature of bacterial chromatin, we must first understand how polyP and DNA interact. Despite the known structural and functional association between these two polyanions *in vivo*, the mechanistic basis of interaction between polyP and DNA have remained poorly understood. A simple Coulombic charge consideration implies that the interactions between two negative point ionic charges, and consequently polyanionic species, is repulsive. The strong repulsive interaction, therefore, must be counteracted by a positive charge for a stable interaction between PolyP and DNA. Peptides, proteins, polyamines, and metals can all drive polyP condensation through phase separation, and likely participate in mediating these interactions^27–29^.

As the protein partners can widely vary across different species and biological systems, we hypothesized that divalent cations could provide a general, system-independent mechanism of interaction between the two polyanionic polymers. Divalent cations are well-known to induce homotypic phase transitions with other polyanions^27,30–32^, notably RNA. The notion that Mg^2+^ may be an important mediator of polyP-DNA interactions *in vivo* is further supported by an early observation where depleting the minimal medium of Mg^2+^ prevented *Aerobacter aerogenes* from making polyP granules^33^. In addition, numerous studies in diverse bacteria have used elemental analysis to show that polyP granules are enriched in divalent cations, including Mg^2+ 8,10,34,35^. Moreover, Mg^2+^ is the most abundant cation in the bacterial cytoplasm, and is believed to be largely bound to nucleic acids. From evolutionary and biophysical perspectives, characterizing the emergent properties of this multicomponent system of polyP-Mg^2+^-DNA is a key starting point to understanding the role of polyP in chromatin structure and function. In this study, we ask: what are the properties of the polyP-Mg^2+^-DNA interface? How does the formation of the multicomponent system affect the organization of DNA? And how does DNA tune the organization and dynamics of polyP condensates?

## RESULTS

### Long polyP undergoes Mg^2+^-driven reentrant phase transitions

As a starting point for our multicomponent system, we first tested the ability of Mg^2+^ to drive phase separation behavior of polydisperse polyP in a length regime that would be expected to be found in bacteria, which can make chains in the 100s to 1000s of orthophosphates in length. Based on our and other previous work on divalent cation-driven RNA/polyanion phase separation^27,30–32,36^, we were interested in understanding the Mg^2+^ concentration dependence and possible non-monotonic characteristics of this process. We therefore charted the Mg^2+^ induced phase separation of long chain polyP (P700-mean: 113 kDa, mode(n_P_): 1000-1300, range: 10kDa - 208kDa) at pH 7.5, as a model *in vitro* system. For these studies, we employed absorption spectroscopy measurements, which can be used to quantify light scattering induced by phase separation, a method that has been previously used for such studies^32,37^. Additionally, confocal fluorescence microscopy was used to visualize the morphologies of the resultant species. For the imaging studies, polyP was labeled with AlexaFluor 647 using a reported procedure^25,38^.

The absorbance data indicate an onset of phase separation around 10 mM Mg^2+^ concentration (Fig 1a). Imaging studies confirmed that the absorbance increase corresponded to formation of spherical droplets that showed facile fusion on the few second timescale, consistent with liquid-like behavior (Fig 1b, SI Movie 1). Bleached regions in polyP-Mg^2+^ condensates reached just under 80% recovery within 50 minutes in fluorescence recovery <u>a</u>fter <u>p</u>hotobleaching (FRAP) experiments (Fig 1c, Fig S1a). Compared to some other protein-RNA systems which can recover within seconds to a few minutes for a similar size of bleached region^39–41^, polyP recovery in polyP-Mg^2+^ condensates is relatively slow.

**Figure 1.**
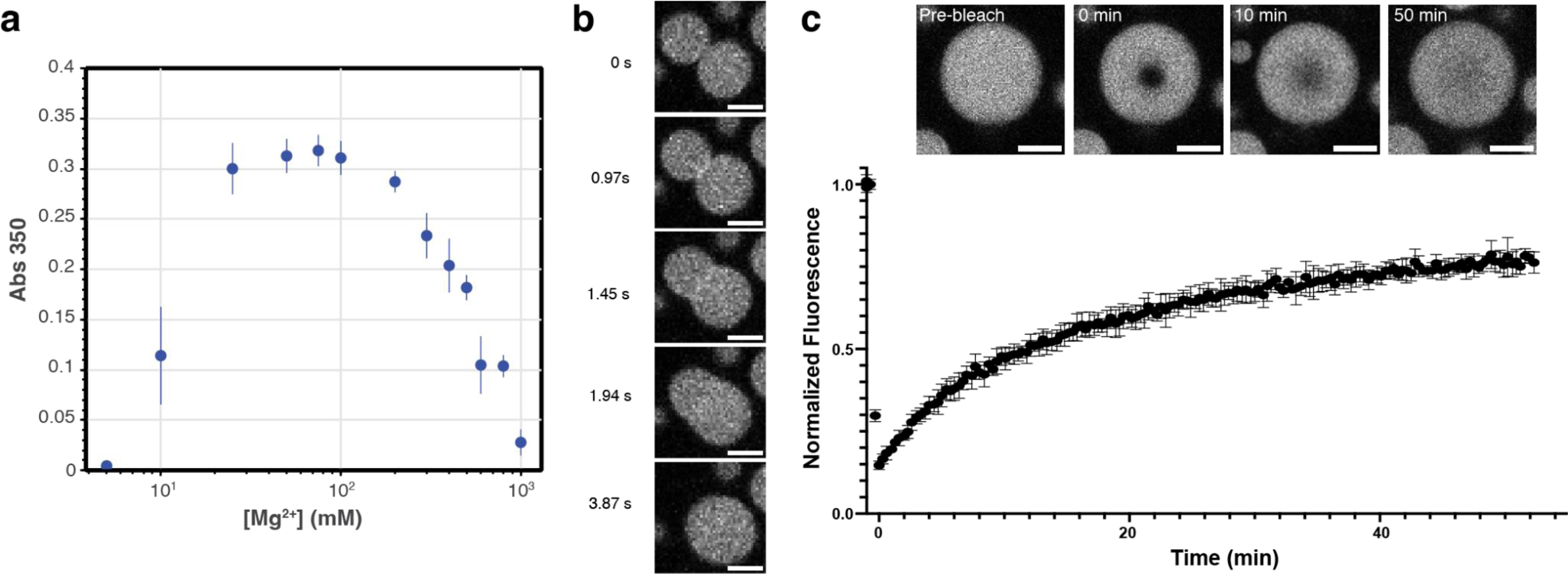
PolyP-Mg^2+^ coacervates exhibit reentrant phase transition and are dynamic. **a** Phase boundary curve for polyP-Mg^2+^ coacervates as determined by the solution turbidity ([polyP] = 1 mg/mL, 50mM HEPES-NaOH, pH 7.5). **b** Representative confocal fluorescence microscopy images of polyP-Mg^2+^ mixtures that correspond to 100mM MgCl_2_ of the phase diagram. Images represent fusion of polyP-Mg^2+^ coacervates ([polyP] = 1 mg/mL, polyP-AF647 = 10% polyP, [Mg^2+^] = 100mM, 50mM HEPES-NaOH, pH 7.5; scale bar = 2µm). A movie showing a larger field of view of droplet fusion is available (SI Movie 1). **c** PolyP-Mg^2+^ coacervates recover to around 80% 50 minutes after photobleaching in Fluorescence Recovery After Photobleaching (FRAP) experiments (d_bleached ROI_ = 1.7µm, d_droplets_ = 8.4-8.5µm, n = 4). Representative images showing recovery at select timepoints are inset (scale bar = 2µm).

The absorbance data also reveal reentrant behavior with a reduction in scattering observed for Mg^2+^ concentrations above 100 mM. This roll-over is similar to the behavior demonstrated previously for RNA-protein and other condensates^37,42,43^. This effect can be attributed to droplet dissolution past the charge-balance region around 100 mM Mg^2+^, where the surface interaction valences of smaller polyP species (single molecules or clusters) are quenched by excess Mg^2+^, thus terminating the network and preventing larger condensate formation. It is noteworthy that complete dissolution is observed at high Mg^2+^ concentration, indicating a lack of residual networking interactions in this reentrant region as observed in some other reentrant systems such as polyrA-Mg^2+ 30,32^. Furthermore, time-series imaging reveals the formation of dynamic vacuolar species during dissolution (Fig S1b&c, Movie 2), similar to reported non-equilibrium dynamics of RNA-peptide complex coacervate systems^37,44^.

Overall, these studies establish the fundamental characteristics of the polyP-Mg^2+^ system for this biologically relevant polyP size range. We observe an onset of phase separation at biologically relevant low mM Mg^2+^ concentrations, along with reentrant behavior and dynamic substructure at higher Mg^2+^ concentrations. Building on these results, we next studied the effects of DNA in the polyP-Mg^2+^-DNA system.

### DNA interacts with polyP-Mg^2+^ droplets, forming shells that display reentrant behavior

We next studied the effects of inclusion of circular double-stranded DNA in the system. Since the observed polyP granule embedding within the bacterial nucleoid may also involve other cellular factors, we aimed to test the intrinsic morphology and physical principles for this simplified DNA-polyP-Mg^2+^ system. Based on prior cellular and *in vitro* work^27,43,45–49^, we could envision several scenarios. These would include partitioning of the DNA into the polyP-Mg^2+^ droplets, or formation of a core-shell architecture, with the interaction of the two polyanions being potentially mediated by Mg^2+^. Other possibilities include formation of DNA-Mg^2+^ condensates that either stay separate from or associate with the polyP-Mg^2+^ condensates. These latter scenarios are less likely given lack of reported evidence of DNA condensation by divalent cations except in a limited set of conditions^50–53^. Additionally, DNA may be excluded from the polyP-Mg^2+^ droplets. Furthermore, we aimed to test if the non-monotonic phase behavior of the polyP-Mg^2+^ components also resulted in modulation of the DNA morphology in the multicomponent system.

First, we used pUC19, a standard, circular 2.7 kb plasmid DNA at a final concentration of 10 µg/mL, labeled with the intercalating dye, YOYO-1. To facilitate droplet visualization and quantification, we first probed the system near the peak concentration of 100 mM Mg^2+^ in the polyP-Mg^2+^ phase transition curve. Experiments were carried out by pre-mixing the two polyanions followed by induction of phase separation by addition of Mg^2+^. Strikingly, upon induction of phase separation by addition of Mg^2+^, we observed that pUC19 DNA formed a shell (Fig 2a, yellow) associated with the surface of the polyP-Mg^2+^ droplets (blue). A 3D construction of confocal fluorescence microscopy images confirms the surface association of DNA across different planes (Fig S2a). Additionally, inspection of the fluorescence intensity profiles across polyP-Mg^2+^ condensates revealed that DNA is preferentially recruited on the condensates’ surface while being relatively depleted from the condensates’ core (Fig 2b). These shells were also visible with 5′ Cy5 end-labeled DNA (Fig S2b). Thus, this DNA forms a shell around the polyP-Mg^2+^ droplets.

**Figure 2.**
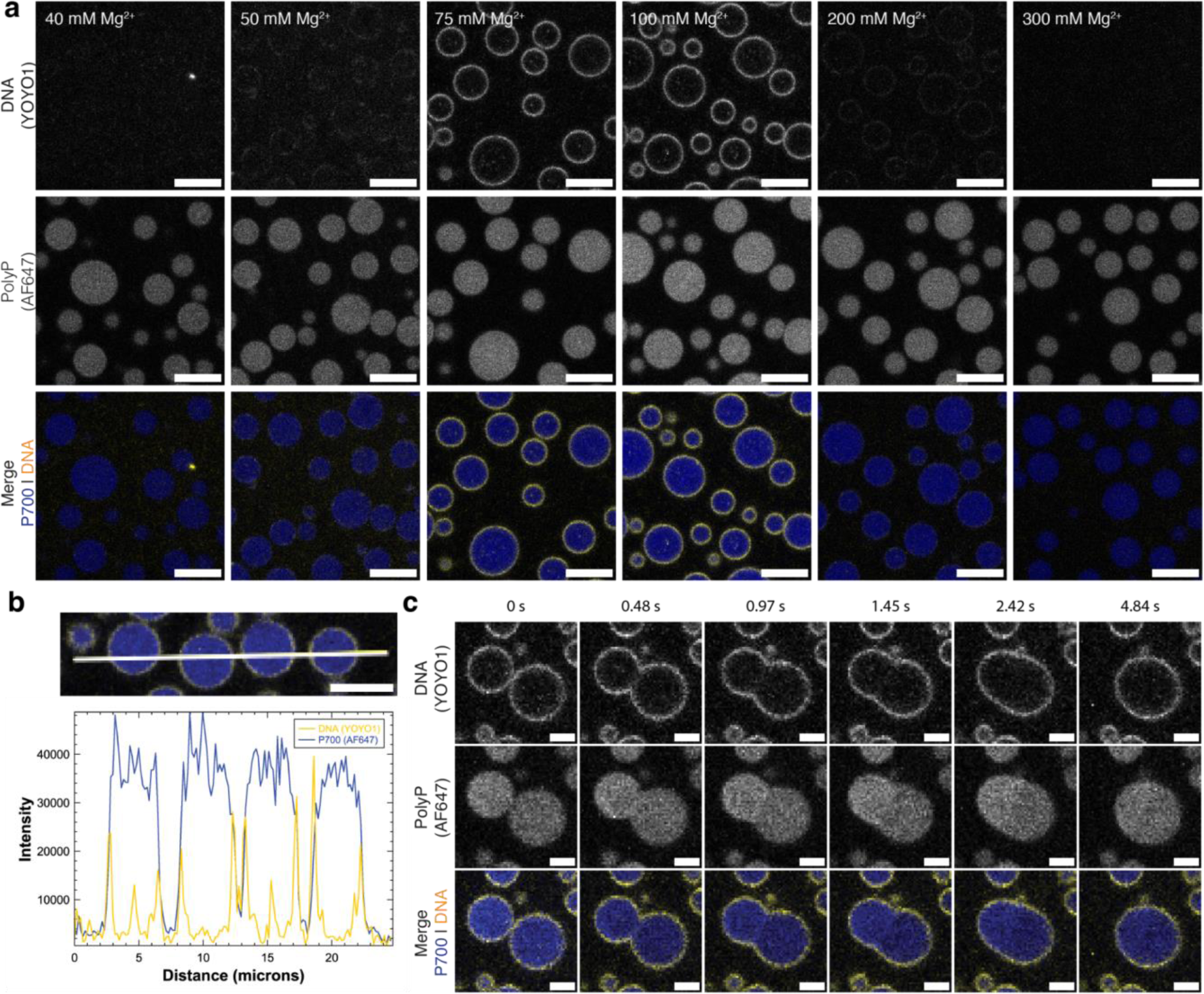
DNA interacts with the surface of PolyP-Mg^2+^ coacervates and forms shells that exhibit reentrant behavior. **a** Confocal fluorescence microscopy of polyP-Mg^2+^ coacervates and pUC19 (2.7kb) plasmid under different MgCl_2_ conditions. DNA forms a shell on the surface of PolyP-Mg^2+^ coacervates within a Mg^2+^ concentration range of 50-200mM. Three channels corresponding to Alexa Fluor 647 (P700), YOYO-1 (DNA) and the merge of these two channels are shown ([polyP] = 1 mg/mL, polyP-AF647 = 10% polyP, 50mM HEPES-NaOH, pH 7.5; scale bar = 5 µm; P700, blue; DNA, yellow). **b** Intensity profiles across PolyP-Mg^2+^-DNA coacervates corresponding to [Mg^2+^]=100mM (other conditions described in panel a) showing the surface localization of DNA (scale bar = 5µm). **c** Confocal fluorescence microscopy images at different time-points of polyP-Mg^2+^-DNA coacervate fusion (for conditions described in b, scale bar = 2 µm). See Fig S2C for the full frame fusion and SI Movie 3 for a wider field of view video.

A key question that arises from these observations is whether shell formation restricts the fusion of polyP-Mg^2+^ droplets. This question is especially relevant given our prior observations that in *P. aeruginosa* under nitrogen starvation conditions, multiple polyP granules are transiently evenly spaced in the nucleoid and do not undergo complete coalescence. An examination of time-lapse images of the system clearly demonstrated rapid fusion of these droplets (Fig 2c, Fig S2c, SI Movie 3). Thus, under these conditions, the circular pUC19 shells do not substantially restrict droplet fusion. We also carried out FRAP experiments to understand molecular mobility in the DNA shells. However, our experiments are not able to clearly distinguish various FRAP contributions from diffusion of DNA and (non-covalently bound) YOYO-1 label. Hence FRAP data for DNA shells are not presented here.

We next asked if the position along the polyP-Mg^2+^ reentrant phase curve would influence the properties of the DNA shell. Since polyP and DNA do not form droplets without Mg^2+^ under our conditions, we hypothesized that DNA interacts with positive charges (Mg^2+^) on the surface of polyP-Mg^2+^ droplets. Based on our prior work on reentrant behavior of RNA-peptide phase separation^37^, we anticipate that there is a charge inversion in the region of the peak in polyP-Mg^2+^ phase separation (Fig 1a), i.e., the surface of the droplets becomes negatively and positively charged in the regions to the left and right of the peak respectively. Therefore, a prediction from the charge-based DNA:polyP-Mg^2+^ droplet interaction model is that shell formation should be reduced in the lower Mg^2+^ concentration region.

To test this model, we carried out a series of imaging experiments, checking for DNA shell formation at different polyP-Mg^2+^ ratios. We observed that in keeping with the interaction model, shell formation was substantially reduced below 50 mM Mg^2+^ (Fig 2a, Fig S2d). Interestingly, shell formation also was not observed above 200 mM Mg^2+^ (Fig 2a, Fig S2d). We can rationalize this latter observation using the same mechanism as we discussed for reentrance in the polyP-Mg^2+^ system. At high Mg^2+^ concentration, the charges on DNA molecules are screened by the excess Mg^2+^, thus reducing the propensity to interact with the droplet surfaces. Although the predominant DNA density appears uniform on the surface, we also observe puncta both on the surface at low Mg^2+^ where shells are less prominent and occasionally within the condensates at Mg^2+^ concentrations where shells form (Fig 2a).

Our results therefore show that pUC19 DNA forms shells at the surface of polyP-Mg^2+^ droplets within a concentration range around the maximum in the reentrant curve in Fig 1a. However, a closer inspection of the images and intensity profiles indicate that the shells may be thin within the resolution limit of our imaging method.

### The condensate interface exhibits distinct morphologies as a function of DNA concentration and length

To determine the morphological features of the DNA shells on the surface of Mg^2+^-polyP condensates, we turned to the higher resolution provided by cryo-electron tomography (cryo-ET). Cryo-ET is a particularly powerful and agnostic approach to determine the structural properties of interfaces at high resolution across a wide range of length scales. We probed how concentration and length, properties that could affect the number of surface contacts, global orientation, and packing dynamics, affect the architecture of DNA at the interface.

Representative tomographic slices of PolyP condensates incubated with different types of DNA are shown in Fig 3a-d, with the associated 3-dimensional renderings shown in panels e-h, respectively (refer to Fig S3a-f for corresponding low magnification images of grids). In the absence of DNA, the interface of polyP-Mg^2+^ condensates exhibits a dense edge (Fig 3b, red arrow; 3h). We also observe a dense edge in the presence of DNA, which could be a combination of PolyP-Mg^2+^ and DNA (Fig 3a-d). To represent this ambiguity the surface rendering displays this feature in yellow (Fig 3e-h). A dense edge has been previously noted for polyP granules *in viv*o in *Acetonema longum* spores and is also visible in several other systems^8,10,54^. With 10µg/mL pUC19 plasmid DNA (2.7kb), we observe distinct filaments protruding from the surface (Fig 3a-d, Fig S4). Condensates formed with 10-fold more DNA (100µg/mL pUC19) exhibit protruding filaments that are both more numerous and extend further from the surface, with some filaments extending more than 100nm from the surface (Fig 3d).

**Figure 3.**
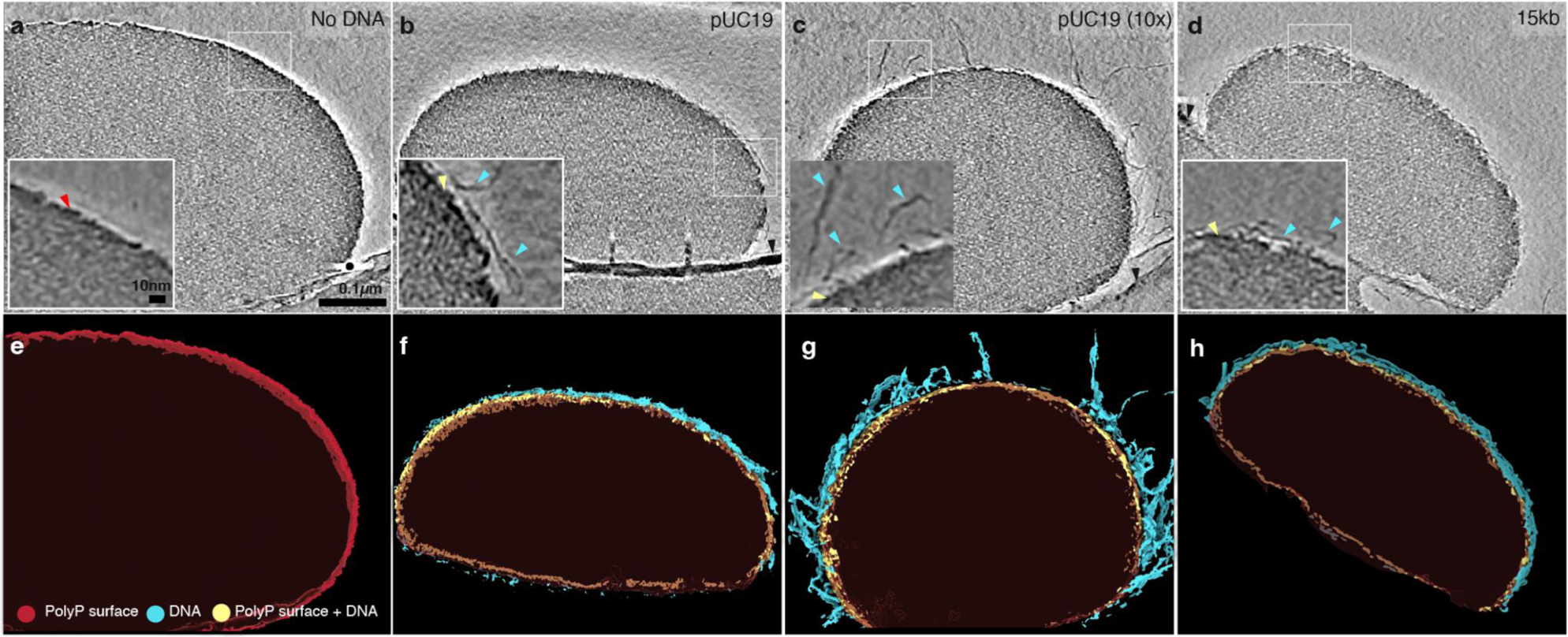
Cryo-electron tomography. (a-d) Representative tomographic slices of PolyP condensates incubated with different types of DNA. Red arrow highlights the dense edge of polyP, cyan arrows highlight DNA and yellow arrows highlight the dense edge+DNA surface (scale bar = 100 nm). (e-h) 3-dimensional renderings of tomograms shown in panels a-d, respectively. The dense edge of PolyP condensate is shown in red, the dense edge+DNA are shown in yellow, and DNAs are shown in cyan.

In the presence of longer, circular DNA (15kb) at 10µg/mL, we observe filaments protruding a similar distance from the surface as with circular pUC19 (Fig 3b, c). Alternative views of the 3D renderings highlighting the different surface textures are available in Fig S5 and SI Movies 4-7.

To determine the effect of DNA concentration and length on the thickness and density of the interface, we performed subtomogram averaging on thousands of randomly selected ∼30nm cubic regions spanning the interface (Fig S6a). We quantified the thickness of the dense edge by drawing eight x-y plane density profiles on the mid-section of the average maps perpendicular to the edge and averaging the thickness values (Fig S6b-g, Fig S7).The measured thicknesses of the dense edge were measured to be 4.6±0.7nm in the absence of DNA, which was not a significantly different upon addition of DNA (Fig S6f). We observe an additional outer layer of intermediate density between background and the dense edge in the presence of 100µg/mL pUC19 DNA (Fig S6d, cyan arrow) which we attribute to the protruding filaments. These findings are consistent with DNA adsorbing to the surface of polyP-Mg^2+^ condensates as a thin shell. The DNA shell’s packing architecture, including length and density of filaments, depends on DNA concentration.

### DNA concentration and length modulate the size of polyP-Mg^2+^ droplets

Our cryo-ET observations provide several key insights into the general structural features of the DNA shells and their dependencies on DNA key parameters. Given the known ability of adsorbed macromolecules to stabilize emulsions and colloids against fusion/aggregation^55,56^, we next returned to fluorescence imaging studies to test whether DNA shells can similarly influence polyP-Mg^2+^ condensate size distributions. This aspect is especially interesting given the transient organization of multiple non-fusing polyP granules in *P. aeruginosa*^10^. We probed the influence of DNA concentration and length in this set of experiments.

#### DNA concentration

We considered several mechanisms that could contribute to the dependence of droplet size on DNA concentration. First, in the case of a thin shell, the total maximum available DNA-polyP interfacial area should be a monotonic function of DNA concentration. Therefore, since the interfacial area of a given volume of polyP-Mg^2+^ condensate will have an inverse dependence on droplet size, higher DNA concentration should result in smaller droplets given that shell formation must overall be energetically favorable. Furthermore, our cryo-ET results revealed that increasing the concentration of pUC19 resulted in a brush-like morphology^57^ of DNA on the droplet surface, which should also result in slowing of droplet fusion and smaller droplets due to the physical/entropic barrier on the droplet surface. Higher partitioning and packing of surface DNA at higher concentrations may also lead to slowing of fusion. Thus, the above thermodynamic and kinetic mechanisms should all result in reductions in droplet size as a function of increasing DNA concentration.

To test this idea, we performed widefield and confocal fluorescence imaging experiments using a series of DNA concentrations ranging from 0 to 100 µg/mL with the same polyP and Mg^2+^ conditions as previously used (SI Figs S8 and S9). Droplets appeared to decrease in size at higher concentrations of DNA (Fig 4a (top/middle), 4b), and we also occasionally observed the appearance of rod-like filaments of micrometer scale (Fig S8 and S9).

**Figure 4.**
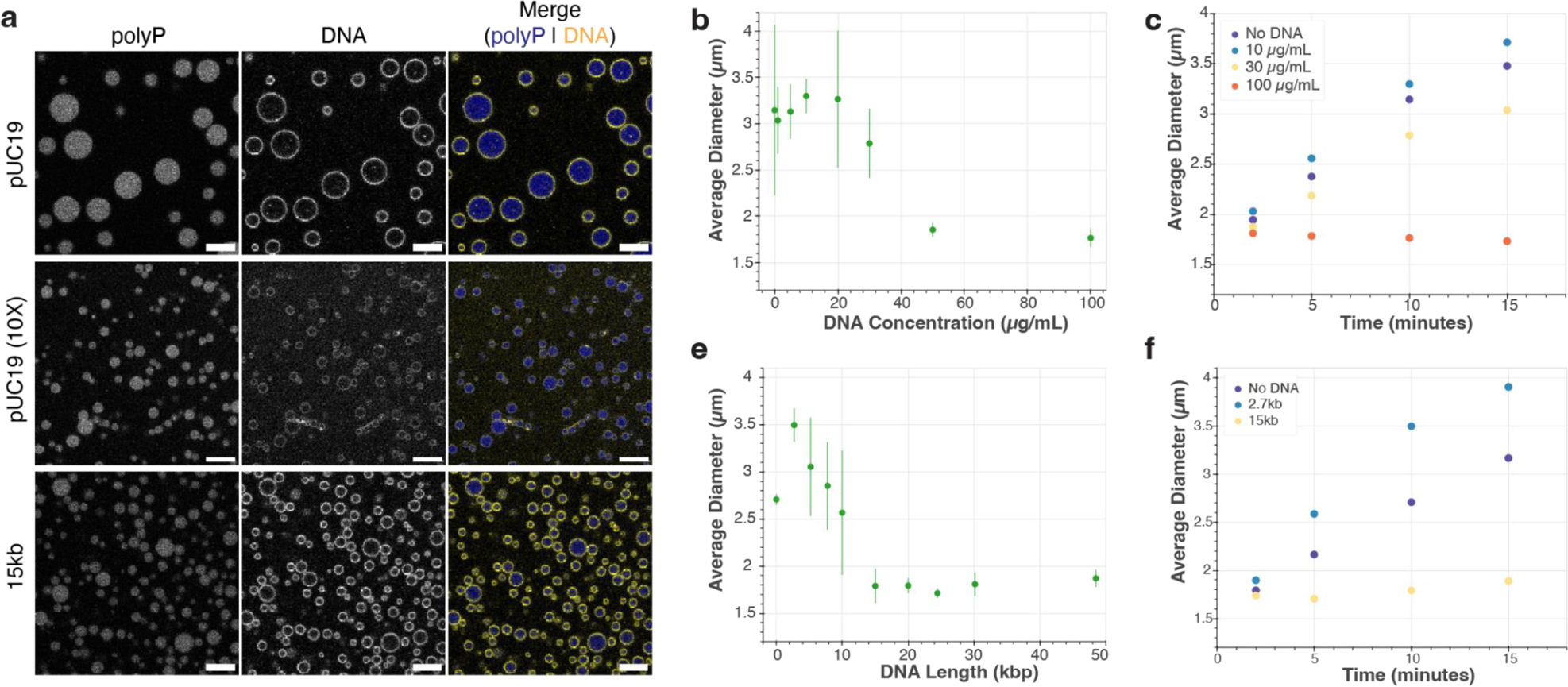
Effect of DNA concentration and length on PolyP-Mg^2+^ size distribution and average droplet size. **a** Representative sample confocal images of polyP-Mg^2+^ droplets given different DNA concentration (top & middle) and length (top & bottom) ([polyP] = 1mg/mL with ∼10% P700-AF647, [DNA] = 10 µg/mL or 100 µg/mL, YOYO1 = 1µM, 50mM HEPES, scale bar = 2µm). Representative confocal images for each of the lengths tested and select concentrations in confocal and widefield are available in SI Figs 8-9, 13-14. **b** Scatter plot showing the average of mean droplet size across three experiments with respect to varied DNA concentrations (error bars = SD of mean diameters of each experiment). At 30 µg/mL DNA, the average droplet size begins to decrease. **c** Scatter plot showing average droplet size as a function of time for three representative DNA concentrations. **d** Scatter plot showing the average of mean droplet size across three experiments with respect to different DNA lengths (error bars = SD of mean diameters of each experiment). DNA length used include circular plasmids of length 2.7kb (pUC19 used for panel a-c), 5kb, 8kb, 10kb, 15kb, 20kb, 24kb, 30kb and commercially available phage DNAs Lambda (49kb) and T4 (166kb). At longer DNA lengths, condensate size decreases. **e** Scatter plot showing average droplet size as a function of time for three representative DNA lengths.

To probe the droplet size distribution quantitatively, we employed a MATLAB-based image analysis routine to analyze the widefield images (refer to Methods section for more details, and also to Fig S10 for representative images of the segmentation step). We then plotted the average of the mean droplet size (Fig 4b) for each replicate distribution as a single statistic to gain insights about our data. The full quantification of the droplet sizes as an empirical cumulative distribution function (ECDF) plot is available in the SI (Fig S11).

Consistent with the mechanisms discussed above, our analyses revealed that increasing the DNA concentration beyond 20 µg/mL indeed led to a decrease in the droplet size (Fig 4b) and left shift of ECDF curves (Fig S11b). Fig 4c right panel shows the time evolution of the average droplet sizes for three representative DNA concentrations. While the average droplet size of polyP-Mg^2+^ droplets for DNA concentrations 10 and 30 µg/mL grows with a net positive slope, the average droplet size of 100 µg/mL remains close to unchanged (slope∼0) with near-overlapping ECDF curves at the four time points studied (Fig 4c and S12) likely indicating arrest in the droplet size growth at high concentrations.

#### DNA length

We next asked whether DNA length can alter droplet size distributions even if the total base-pair concentration in solution remains constant. It is well known from the polymer physics field that polymer length can intrinsically affect phase separation propensity, often discussed in terms of polyvalency in the condensate literature^58–61^. Previous studies on DNA condensation as well as phase separation demonstrate DNA-length dependent properties^53,62^. In the present context, rearrangements of the surface bound DNA may be increasingly slower as the length increases due to increased avidity or entanglement effects. Since this DNA rearrangement is likely important in droplet fusion kinetics, we hypothesized that shell formation with longer DNA could result in slower droplet fusion and a consequent smaller droplet size.

To test this model, we probed the length-dependence of DNA on droplet formation with polyP-Mg^2+.^ We compared the effects of a range of DNA sizes by using 10μg/mL circular plasmids of length ∼ 2.7kb (pUC19 used thus far), 5kb, 8kb, 10kb, 15kb, 20kb, 24kb, 30kb and commercially available linear phage DNAs Lambda (49kb) and T4 (166kb) (refer to Table 1 for exact DNA lengths and additional details). We chose this range of DNA lengths to span a range from below to above the size of bacterial plectonemes (∼10-15kb): 10kb based on EM, simulations, and gene expression microarray in *E. coli* by and 15kb based on Hi-C and modeling in *C. crescentus*^63,64^. As with the concentration based experiments, we used widefield fluorescence images coupled with MATLAB to quantify their condensate size distributions (Fig S13) and confocal imaging to confirm the presence of 3D shells (Fig 4a (top/bottom), Fig S14-S15). The resulting average size and time-dependence data are shown in Fig 4d-e (also see ECDFs, Fig S16 & S17). Consistent with the above hypothesis, our experiments revealed that increasing the DNA length in the range of 2.7 to 15kb shifted the size distribution of polyP-Mg^2+^ droplets to a smaller size (Fig 4c; left-shifted ECDFs in SI Fig S16), also reflected in the time-dependence (Fig 4e). However, there were some deviations from this trend. First, the size roughly leveled off between 15 and 30 kb, which could be due to substantial growth arrest or because the distribution is clustered close to the resolution limit of our analysis. Nonetheless, fusion can still be observed with the longer 15kb DNA (SI Movie 8). Curiously, T4 DNA exhibits a wider and seemingly anomalous droplet size distribution, which is also reflected in a larger average droplet size and small but positive growth compared to the 15 and 30 kb range (Fig S17); the droplets also tend to cluster together, occasionally moving as a grouped unit (SI Movie 9).

**T1.** Table of plasmids used in our study.

| Plasmid Name | Source | Plasmid Size (bp) | Growth strains |
| --- | --- | --- | --- |
| 2.7kb plasmid | pUC19, NEB Catalog# N3041S | 2686 | DH5alpha |
| 5kb plasmid | Addgene Catalog# 49795 (Xu et al, 2012)<sup>105</sup> | 5203 | DH5alpha |
| 8kb plasmid | Racki Lab plasmid# LR562 | 7768 | DH5alpha |
| 10kb plasmid | Racki Lab plasmid# LR556 | 9988 | DH5alpha |
| 15kb plasmid | Addgene Catalog# 111444 (Vyas et al, 2018)<sup>106</sup> | 15014 | DH5alpha |
| 20kb plasmid | Addgene Catalog# 29036 (Portales-Casamer, 2010)<sup>107</sup> | 20005 | DH5alpha |
| 24kb plasmid | Addgene Catalog #136828 (Lukinavicius et al, 2013)<sup>108</sup> | 24445 | DH5alpha |
| 30kb plasmid | Addgene Catalog# 117760 (Shepherd et al, 2017)<sup>109</sup> | 30152 | BLR (F- ompT hsdSB(rB-mB-)gal dcm (DE3) Δ(srl-recA)306::Tn10 (TetR)) |
| Lambda DNA | NEB Catalog# N3011L | 48502 | - |
| T4 DNA | T4 GT7 DNA, Catalog# 318-03971, FUJIFILM Wako Chemicals USA | 166000 | - |

Overall these length and concentration observations are particularly striking, given the substantial effects observed even at DNA phosphate concentrations 2-3 orders-of-magnitude lower than the PolyP polyphosphate concentration.

## DISCUSSION

Biomolecular condensates have emerged as a key structural feature of both eukaryotic and, more recently, bacterial chromatin^25,65–73^. Diverse partners can drive chromatin condensate formation, but the role of polyphosphate, a universal and ancient inorganic polymer, has been largely overlooked in chromatin biology. We hypothesize that polyP condensates are a fundamental feature of bacterial chromatin, and likely important for chromatin structure and function in all three domains of life.

Empirically, magnesium has been shown to be the dominant cation in bacterial polyP condensates and divalent cations can drive polyP condensate formation, as they do with RNA. Given the critical role of magnesium in nucleic acid structure and function, and the longstanding observation that polyP condensates are embedded in the bacterial nucleoid in diverse species, in this study we have established a fundamental interaction between DNA and polyP mediated by magnesium that determine the properties of these condensates. We discovered that DNA associates with the surface of polyP-Mg^2+^ coacervates. This surface association both affects the morphology of the DNA and tunes the size of the condensates in a manner dependent on DNA properties.

### PolyP-Mg^2+^ coacervation

In our study, we found that interactions between long chain polyphosphate, relevant to bacterial physiology, and Mg^2+^ can result in the formation of coacervates. The formation of coacervates of longer length PolyP in presence of Mg^2+^ is consistent with the larger body of polyP-Mg^2+^ coacervation work in the context of phosphate glasses and more recently in the context of RNA interactions and condensation^74^. Our observed onset of condensation in this system (∼10 mM Mg^2+^) is substantially lower than reported thresholds of Mg^2+^-induced phase separation of long polyU RNA and similar to that of short polyA RNA in the absence of crowding agents^27,30,32^. Additionally, while relatively rapid fusion resulted in spherical droplets, our FRAP results showed that diffusion and mixing within the resultant droplets were slow, qualitatively similar to previous observations in chromatin^75^ and the much slower internal rearrangement of polyrA in Mg^2+^-induced condensation^32^. Given these observations, it is worth noting that the condensates studied in this work could be considered as network fluids that are expected to exhibit viscoelastic characteristics^28,58,61,76–78^, an important direction for future work. Given the similarities of our system with other homotypic coacervates of RNA and divalent cations^32,36^, we predicted that the system would only result in coacervation in a window of relative polyP-Mg^2+^ concentrations around the charge-balance region. Our demonstration of precisely this type of reentrant behavior highlights the importance of charge-based interactions in mediating networking in these coacervates. Motivated by the previous observations in RNA-peptide systems^37,42,44^, we also tested and verified that a jump of Mg^2+^ concentration can lead to the formation of dynamic, non-equilibrium vacuole-structures, the *in vivo* implications of which remain to be determined.

We also observe the presence of a dense edge in our cryo-ET images (Fig 3a-d), which appears even in the absence of DNA. This is particularly interesting given that a dense edge has also been previously noted for polyP granules *in viv*o in *Acetonema longum* spores^8^. While the dense edge has been hypothesized to be the product of proteins gathering on the surface, it is interesting that a similar feature can be recapitulated *in vitro* in a system containing only polyP and Mg^2+^. We speculate that the dense edge may be an outcome of differential hydration of Mg^2+^ at the surface compared to the droplet interior, and could be similar to differences in hydration, ion concentration, and binding observed in polyP-Ca^2+^ systems^79^.

### DNA-association with Mg^2+^-polyP condensate surfaces

Our studies have also revealed that DNAs are preferentially recruited on the condensates’ surface while being relatively depleted from the condensate core. The association of DNA with the PolyP-Mg^2+^ surface presumably arises from favorable interactions between the negatively charged phosphate groups on the backbone of DNA and Mg^2+^ at the surface of polyP-Mg^2+^ coacervates. Such a model would also be consistent with differential hydration of Mg^2+^ inside and at the surface of PolyP coacervates discussed in the previous section, and could lead to the emergence of unique surface properties relative to the internal condensate environment. A charge-based interaction is consistent with our observations of the reentrant nature of the DNA shells which form under a relatively narrow range of Mg^2+^ concentrations, where we expect both the surface to be positively charged/near-neutral and the divalent cation concentrations to be within a regime that does not screen charge-based interactions.

While higher-order core-shell architectures have been observed both in cells/*in vivo* and recapitulated *in vitro*^27,39,45–48,80–84^, there are some notable differences between these multiphase condensate systems and our own. First, in contrast with many previous studies with more comparable concentrations of the different biopolymers, we studied a region of concentration space where DNA phosphate concentrations were generally more than two orders of magnitude lower than those of polyP-Mg^2+^ (for the majority of experiments, ∼15 *μ*M DNA phosphate vs ∼10 mM polyP phosphate and >10 mM Mg^2+^). Our results demonstrate that even such extremely small relative concentrations of DNA can exert substantial control on certain properties of polyP-Mg^2+^ condensates which has potential implications for other cellular condensates where minor or undetected components could be important for biological regulation and function. Another important contrast with many other described core-shell systems is the lack of DNA condensation in similar Mg^2+^ concentration regimes in the absence of PolyP. Indeed, to the best of our knowledge, divalent cations (like Mg^2+^ and other alkaline earth metal ions) are not known to condense dsDNA in dilute, bulk solution alone and require additional special conditions like addition of PEG to induce DNA condensation (termed PSI-condensation) or change of solvent conditions (such as changes in dielectric constant) ^50–52^. On the other hand, similar to previously discussed mechanistic understanding for multiphasic core-shell condensates^45–48^, it is likely that an overall reduction of the interfacial energetic cost is one driving force for DNA shell formation in our polyP-Mg^2+^-DNA system.

While we cannot rule out the possibility that the multicomponent system here is a form of the multiphase condensates described above, it is tempting to speculate that PolyP-Mg^2+^ induces the adsorption and subsequent condensation of DNA on its surface. We note potentially related observations of adsorption and formation of shell-like structures in Pickering emulsions and some RNA-based condensates^85^. Such surface induced adsorption and condensation would be consistent with previous work showing DNA adsorption/condensation on cationic and zwitterionic lipid surfaces^51,86–90^. Interestingly, for the studied zwitterionic systems, these surface-based interactions appear to be mediated by the divalent counterion Mg^2+^. Given the thin nature of the DNA shells observed in our work (Figs 2 and 3), the presumed surface charge dependence of the interaction, and the notable absence of a separate DNA-Mg^2+^ dense phase, our multicomponent polyP-Mg^2+^-DNA system thus potentially represents a novel system for studying 2D-DNA condensation, adding to and complementing previously studied DNA-lipid systems.

### DNA tuning of droplet growth

We rationalized the differences in droplet size from varied DNA concentration and length to originate from a combination of both thermodynamic and kinetic driving forces. Indeed thermodynamic arguments might explain some of the DNA morphology we observe at the interface and the emergence of shells. However, many of our quantitative observations cannot be explained by thermodynamics alone and instead suggest that kinetic factors could play a significant role in controlling droplet growth.

For the DNA concentration dependence, simple consideration of energetic stabilization by shell formation would be consistent with higher DNA concentration correlating with smaller droplets, since the system would try and maximize the DNA-polyP interfacial area. However, that model assumes similar shell morphology for the different DNA concentrations. In contrast, we clearly observe a much more extended brush-like DNA morphology at the 10X DNA concentration, consistent with a physical barrier for fusion and growth. Overall, we therefore conclude that a combination of thermodynamic and kinetic contributions give rise to our observed concentration dependence of droplet size. As a related note, naturally occurring polymer brushes are important in attenuating interactions of large macromolecular assemblies in a variety of biological systems^91,92^. And polymer brushes have been harnessed in diverse engineering and industrial applications to prevent flocculation of particles^93^.

Similarly, we attribute trends observed from DNA length variation to be a consequence of kinetic and thermodynamic contributions. Using the same simple thermodynamic consideration, it could be expected that maintaining the same base pair concentration of DNA could result in minimal changes to droplet size given the constant potential for contacts with the polyP-Mg^2+^ condensate surface. However, we observe a clear dependence on DNA length. An interesting consideration is that kinetically driven differences such as ease of rearrangement, entanglement or jamming^58,86,94,95^ that scale with DNA length could play a role. Properties that scale non-linearly with DNA length including unequal numbers of effectively available contacts due to constraints in DNA bending from supercoiling could also contribute. Moreover, the dependence could also be a product of several thermodynamically driven differences such as partitioning/binding affinity to the surface favoring longer DNA.

### Open questions and functional implications of DNA shells

Our results demonstrate that the DNA shell architecture can affect polyP coacervation, but it is also possible that the properties of DNA are altered in functionally important ways as a consequence of this association. Indeed surface association can dramatically alter the properties of polymers. This is well established with polymer brushes, where increased packing density drives polymer extension through repulsive interactions or entropic effects^92,93^. Interestingly, in the case of polyelectrolyte brushes, multivalent cations can oppose this effect, leading to more collapsed configurations^92,93^. Magnesium bridging interactions are thought to enable DNA to pack more densely when spatially confined, including confinement to a 2D surface^52^. Thus packing density and divalent cation partitioning at the interface of polyP condensates may dynamically tune DNA compaction. Given that polyP synthesis is upregulated during growth arrest, condensate formation may be a mechanism to regulate local DNA compaction. Additionally, our striking observation of reentrance in shell formation suggests a potential avenue for cellular regulation, as has been invoked in the case of RNA-protein reentrant behavior^42,43,96^. Lastly, since DNA supercoiling can affect many processes, notably transcription, and DNA adsorption of charged surfaces can alter supercoiling^97–99^, it is intriguing to speculate that *in vivo* interaction with polyP granules could module DNA supercoiling and associated function locally and more globally^100^.

In this work, we have explored the surprisingly complex and tunable Mg^2+^-mediated condensation behavior of two polyanions with broad relevance in biology and other fields (Fig 5). Of course *in vivo* other factors, including chromatin binding proteins, participate in mediating the interaction between DNA and polyP, as has been shown for the NAP Hfq^25^. Such interactions may act to bring specific DNA loci to the surface and further tune the conformational state of the DNA. NAPs may also modulate the partition of DNA between the surface and the interior, change the properties of the condensates, and provide additional interactions that substitute for and compete with interactions with cations. For example, the histone H1-like protein AlgP in *P. aeruginosa*, which has a +55 charge at neutral pH, localizes to the granules and alters their consolidation dynamics^26^. Furthermore, our results indicate that other polyvalent cationic species, including other divalent cations, whose concentrations can vary in response to both extracellular and intracellular cues, can likely mediate and tune polyP granule formation, as well as their interactions with DNA and materials properties, and will be an exciting avenue for future investigation. From a biophysical perspective, it would be interesting to expand this system to include both chromatin proteins such as AlgP and other well-represented biological polyanions, specifically single-stranded RNA and DNA (Fig 5). This future direction is particularly relevant given that these polyanions are widely represented in cellular condensates, including ones involved in transcription and RNA regulation^85,101,102^. Additionally, such single-stranded systems can add a more complex conformational landscape than duplex DNA, another interesting feature with potential broad biological, chemical and prebiotic relevance.

**Figure 5.**
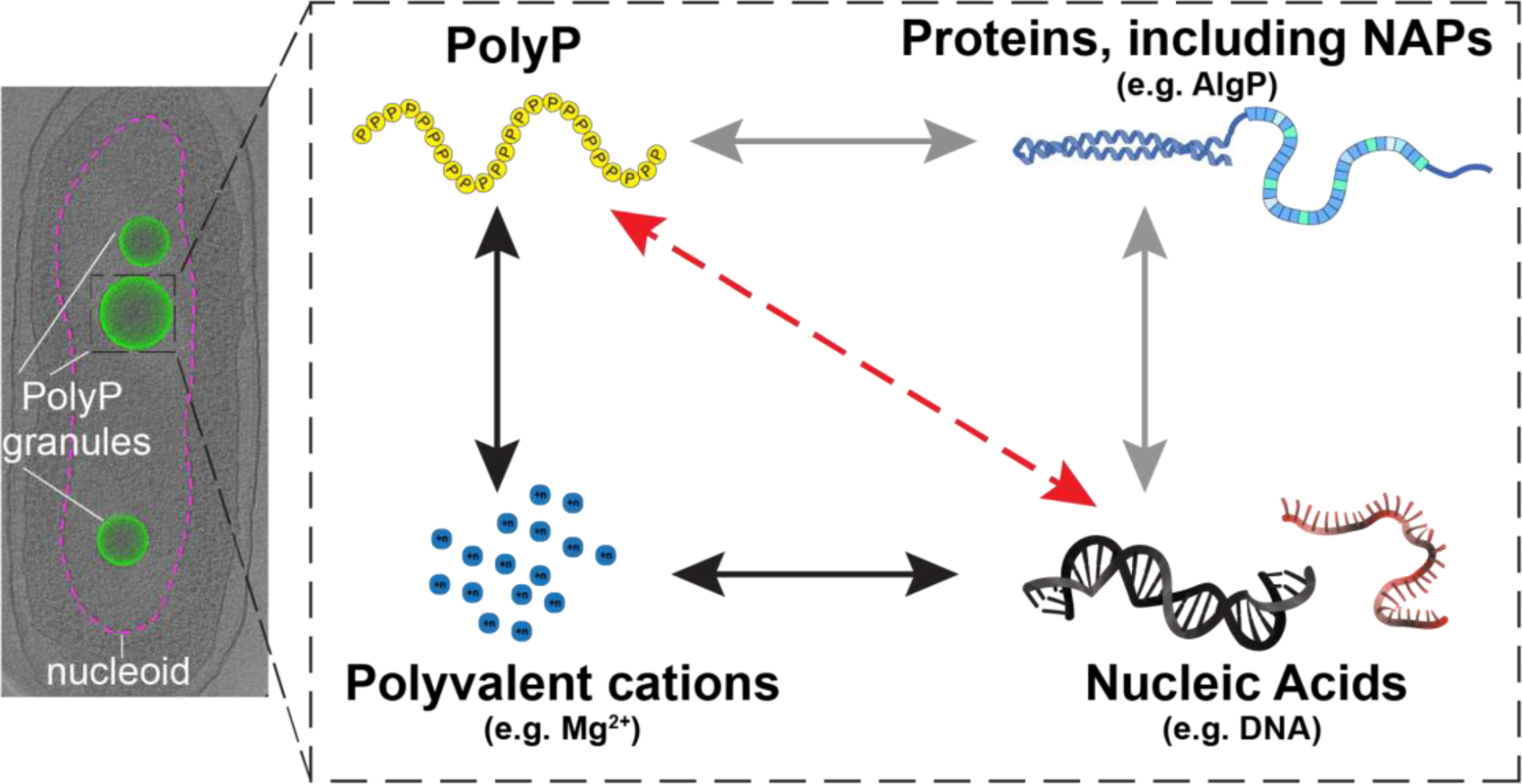
A framework for understanding polyP-chromatin interactions. Left: Cryo-ET of nitrogen-starved *P. aeruginosa* cells with nucleoid region (ribosome depleted) delineated with dashed magenta line, polyphosphate granules shown as green spheres (image: Racki et al., 2017^10^). Right: In this study we have developed a three-component polyP-Mg^2+^-DNA system (interactions represented by black arrows) which is a fundamental physicochemical interaction unit underlying the functional coupling between polyP granules and chromatin in cells. Red dashed arrow represents repulsive interactions between the polyanions, and polyvalent cationic species and proteinaceous partners, including NAPs, represent factors that mediate this interaction. Our results highlight the tunable nature of this minimal system, showing that DNA interacts with and forms reentrant shells around polyP-Mg^2+^ condensates, and modulates condensate size in a DNA length and concentration dependent manner. Future studies building on this framework to include relevant proteins such as nucleoid associated proteins (NAPs) known to associate with polyP in vivo (Hfq and AlgP, for example) are needed to understand how polyP affects chromatin structure and function in cells (gray arrows).

## MATERIALS & METHODS

### Reagents and Stocks

Long chain Polyphosphate P700 was obtained from Kerfast, Inc. (EUI002). This high polyphosphate is heterogeneous in size, with approximate polymer lengths ranging from ∼200-1,300 phosphate units; modal size is about 700 phosphate units. We prepared 100mg/mL stocks of P700 in water and stored them at −80°C for long term storage. The 100 mg/mL P700 stocks were used to further prepare sub-stocks of P700 at 10 mg/mL which were stored at −20°C. These sub-stocks were used for experiments. Magnesium chloride was obtained in dried form (M9272-100) as well as 1M MgCl_2_ solution (M1028-100) from Sigma. HEPES solid powder (H3375-100) was obtained from and 1M stock was prepared in deionized water with the pH adjusted to 7.5 by addition of 10N NaOH(306576-100). The stock was stored at 4°C for long term storage. Aliquots of DNA labeling dye YOYO-1 (ThermoFisher, Y3601) were stored at −20°C.

### PolyP Labeling

We adopted a previously developed polyP end labeling protocol with minor modifications^38,103^. Briefly, a reaction of P700 with EDAC and AF647 cadaverine (Sigma, A30679) was set up in the MOPS buffer, pH 8.0 in dark at 37°C in a 1.5mL eppendorf tube. The final concentration of P700, EDAC, AF647 cadaverine and the buffer in the reaction mixture were 100μM (defined in terms of phosphate ends), 150mM, 2mM (20X excess) and 100mM MOPS, pH 8.0 respectively. The eppendorf tube was agitated occasionally (every 10-15 min). At the end of 1h incubation at 37C, the reaction was stopped by placing the eppendorf tube on ice and centrifuged briefly to remove any condensation from the top of the tube. Next, excess dye removal was carried out using spin desalting columns. To remove excess dye and buffer exchange (into water), we employed three consecutive 0.5mL Zeba™ Spin Desalting Columns (7K MWCO) and followed manufacturer’s guidelines.

### DNA plasmid preparation

To cover a range of DNA sizes, we used plasmids that were in our laboratory as well as commercially available DNA like Lambda-DNA and T4. The in-house plasmid preparation was carried out following the manufacturer’s protocol (Qiagen Midi-kits) and eluted in deionized water. Lambda and T4 DNA were dialyzed from the TE buffer into deionized water using Pur-A-Lyzer Mini Dialysis Kit. The DNA stocks were maintained at −20°C and thawed on ice prior to the experiments. The plasmids used for Cryo-ET were purified using phenol-chloroform extraction^104^. The stocks were stored at −20°C.

### Cy5 end labeling of DNA

Plasmid pUC19 was linearized by using restriction enzymes HindIII (NEB) and XbaI (NEB) and purified using NEB minprep kit and ligated with a Cy5 oligo following a previous protocol. Briefly, a 15 times excess of Cy5 labeled primer (pRRC11_56bp_Cy5; /5Cy5/acggccagtgaattcgagctcggtacgatcctctagagtcgacctgcaggcatgca) was ligated to linearized pUC19 using T4 ligase in an overnight ligation reaction at room temperature. The excess oligos were removed from ligated DNA using CHROMA SPIN columns and purified DNA was used directly for microscopy experiments.

### Sample Preparation

#### Absorbance measurements of PolyP-Mg^2+^

Absorbance measurements were carried out with only unlabeled polyP. Sample absorbance was measured 15-20s after droplet induction, with absorbance reported at 350nm (Nanodrop). To ensure proper mixing, the solution was pipetted up and down 3-4 times measurement on the Nanodrop. Final concentration of the system: polyP: 1 mg/mL (unlabeled), 50mM HEPES-NaOH, pH 7.5, [MgCl_2_]: 0-1000 mM. The time of induction of droplets by addition of MgCl_2_ was used as a reference of t=0min for all of our experiments.

#### PolyP-Mg^2+^ condensates with DNA of different lengths

*DNA Length experiments*: Unlabeled P700 and P700-AF647 were thawed from −80°C on ice prior to each experiment. A 10X master mix was prepared by adding 100mg/mL of unlabeled P700 and purified AF647-labeled P700 (termed 10X polyP mixture henceforth). DNA stocks were removed from −20°C and allowed to thaw on ice at room temperature prior to use in the experiments. Buffer (HEPES-NaOH pH 7.5) was added to DNA in a PCR tube followed by incubation with YOYO-1 dye for 7-8 min. After incubation of DNA with YOYO-1, 10X polyP master mix was added to this solution and droplet induction carried out by mixing an equal volume of appropriate 2X MgCl_2_ solution. Typically 3-4 fields of views were acquired per time point (t= 2, 5, 10 and 15 min) for three experiments carried out on different days using widefield microscopy. To ensure proper mixing, the solution was pipetted up and down 3-4 times before being introduced to the glass chamber for observation under the microscope (confocal/widefield). Final concentration: PolyP: 1 mg/mL unlabeled, with ∼10% labeled P700-AF647, 50mM HEPES-NaOH, pH 7.5, [MgCl_2_]: 0-300mM, DNA concentration: 10 μg/mL, YOYO-1: 1 μM. Note: the control experiment for the ‘No DNA’ case had DNA replaced with water and had a final [YOYO-1] = 1 μM in the solution. Note: All droplets were observed at room temperature. The time at which the MgCl_2_ solution was added to induce droplet formation was used as t=0 min reference in all our studies.

#### PolyP-Mg^2+^ condensates with different DNA concentrations

Concentrated pUC19 stock was removed from −20°C and thawed on ice at room temperature prior to the experiment. The concentrated stock was then used to prepare dilutions of DNA stocks for each experiment. Buffer (HEPES-NaOH pH 7.5) was added to thawed DNA in a PCR tube followed by incubation for 5-7 min. 10X polyP master mix was added to the DNA-buffer solution and droplet induction carried out by mixing MgCl_2_ solution as noted previously. Typically 3-4 fields of views were acquired per time point (t= 2, 5, 10 and 15 min) for three experiments carried out on different days using widefield microscopy. To ensure proper mixing, the solution was pipetted up and down 3-4 times before being introduced to the glass chamber for observation under the microscope (confocal/widefield). Final concentration: polyP: 1 mg/mL unlabeled, with ∼10% labeled P700-AF647, 50mM HEPES-NaOH, pH 7.5, [MgCl_2_]: 0-300 mM, DNA concentration: 10 μg/mL. Note: We controlled for the addition of variable YOYO-1 corresponding to DNA concentration in these experiments by completely omitting the addition of YOYO-1, including the control case of No DNA.

### Microscopy and Analysis

#### Confocal Microscopy

Confocal images were recorded on a Zeiss LSM 780 laser scanning confocal microscope. Samples were imaged at room temperature using a 100× oil immersion objective (Plan-Apochromat 100×/ NA 1.40 Oil DIC M27) at a 16 bit depth with pixel size between 0.17 and 0.08 μm. DNA, through YOYO1 labeling, was imaged using an Argon laser set at 20% laser power, which excited at 458 nm. The detection range for the YOYO1 channel was set from 487-561 nm. Detector gain was adjusted to 800 and an offset of 450 was applied to reduce undersaturated pixels. PolyP was detected through P700 labeled with Alexa Fluor 647. A HeNe laser set at 40% laser power was applied, exciting at 633 nm. Detection range was set to 637-755 nm with a gain of 800 and an offset of 300. The imaging settings were held constant for all confocal images except for the polyP-Mg^2+^ only images and movies used in Fig 1 and SI Movie 1, where intensity of the HeNe 633nm laser at the same detection range was set to 5% and the pinhole for the singular laser was adjusted to 105.5 (or 1AU).

Z-stacks were collected at 2, 5, 10, and 15 minutes for samples at different locations for each time point. The frames were separated in z by 0.37μm, except when otherwise noted. For movies acquired through confocal imaging, frames were collected with no fixed delay resulting in a temporal frame separation of ∼484 ms unless otherwise noted.

Images were imported into FIJI^110^ where timestamps and scale bars were added. Some frames were cropped to highlight particular features (e.g., single droplet fusion) or for scaling. No other corrections to the images (e.g. brightness and contrast) were made for all non-FRAP images. Orthoviews and 3D orthosliced views were generated using Imaris Software (RRID:SCR_007370).

#### Fluorescence Recovery After Photobleaching (FRAP)

FRAP experiments of polyP-Mg^2+^ condensates were conducted using the Zeiss LSM 780 laser scanning confocal microscope conditions as noted above.

Samples were prepared by adding an equal volume of MgCl_2_ solution to P700 labeled with ∼10% P700-AF647 in HEPES buffer such that final concentrations were 1mg/mL polyP, 100mM MgCl_2_, 50mM HEPES, pH 7.5. Condensates were allowed to coalesce and fuse for 35-45 minutes, after which a condensate with a diameter around 8.5 μm was selected. The offset for the z-plane was calibrated for reflection autofocus.

Each experimental run collected images at three time points before subsequently initiating a bleaching protocol. Bleaching consisted of two rounds of 15 iterative pulses over a circular region at the center of the droplet with diameter of 1.6μm at 100% HeNe laser power set to a reduced scan speed (pixel dwell time: 12μs). Following bleaching, images were collected in 20s intervals for 52 minutes with reflection autofocus being applied every 15 scans or roughly every 5 minutes.

To correct for drift in the xy dimension over the 52 minutes, images were processed in FIJI where the StackReg plugin^111^ translation transformation was applied to a cropped frame of the bleached droplet. A circular region equivalent to the bleached ROI size was placed at the bleaching area and measured using FIJI’s measure function. Two ROIs equivalent in size and shape to the bleached ROI were used as references for photobleaching in condensates of around the same size as the bleached condensate and were measured in FIJI. Time was adjusted to be zero immediately after the bleach by subtracting the time of the fifth scan from all times.

Data from the transformed bleached ROI corresponding to different time points were double normalized following the equation:

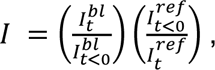

where 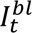 is the average pixel intensity of the bleached ROI at time t, 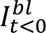 is the average of the three pre-bleach ROI mean pixel intensity, and 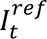 and 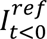 are the corresponding averages for the two reference ROIs.

#### Widefield microscopy

Microscopy images for image analysis were collected using Nikon Ti2-E inverted microscope with perfect focussing system (PFS) and a 100X oil immersion objective (Plan apochromat phase contrast, N.A. 1.45) at a 16 bit depth with pixel size of ∼ 0.11 μm. For brightfield, a white LED, and for fluorescence, the Spectra X Light Engine with a 470nm LED (Lumencor) were used as illumination sources. The camera used for imaging was Prime 95B sCMOS (Photometrics). Image acquisition was controlled using Nikon Elements. Following parameters were typically used: For phase contrast: 10% light intensity, 100ms exposure time, gain = 1.0. For YOYO1 imaging: 5% light intensity from the 470nm LED, 30-100ms exposure time and a GFP filter cube (466/40nm excitation filter, 525/50 nm emission filter, 495nm dichroic mirror, Semrock), gain = 1.0. For AF647 imaging: 10% light intensity from the 640nm LED, 30-100ms exposure time and a quad LED-DA/FI/TR/Cy5-A filter cube(DAPI / FITC / TRITC / Cy5 - Full Multiband Quad Lumencor C19446). For Cy5 imaging: settings similar to AF647 imaging, with the exception of 50% light intensity.

Representative widefield images in the SI were processed using FIJI. Adjustments were made to brightness/contrast by setting the minimum and maximum intensity value to the overall observed min and max values based on the set’s histograms and applying that range equally to all comparable figures. For polyP visualized with the 640 channel, the min and max were set to 1616 and 15601 respectively, while values of 904 and 23335 were used for DNA shells visualized with the 488 channel. The Cy5-end labeled DNA was rescaled to 4231 and 10278. In Fig S6, the brightfield image min was set to 4438 and the max at 34318. No other image intensity modifications were made.

#### Size quantification

Images of condensates from different fields of views and experimental conditions at time points corresponding to t=2, 5, 10 and 15 min were acquired by widefield microscopy. Channel corresponding to 640 (P700-AF647) was used for segmentation and droplet size quantifaction. Custom MATLAB scripts were used for image analysis. Briefly, pre-processing was performed using in-built matlab function *imadjust* and *imclearborder*. Function *imadjust* maps the intensity values in grayscale image to increase the contrast of the output image and *imclearborder* function was used to exclude the droplets at the edge in any given field. MATLAB function *imfindcircles* that employs circular Hough transform was used to find circles in the images. Given the limited accuracy of *imfindcircles* when the value of radius (or *rmin*) is less than or equal to 5, a default *rmin* of 6 was used for all of our analysis. Note: the use of *rmin* sets a minimum radius of droplet detection as 0.66 μm (or diameter 1.32 μm). A default value of parameters *rmax*=90 and *sensitivity*=0.85 were used for *imfindcircles* and adjusted as needed for each field of view to capture the most accurate size distribution using manual visual inspection. The codes were able to accurately capture size distribution for larger sizes; we would, however, like to note that the codes were not able to capture droplets with sizes less than *rmin* 0.66μm and sets a lower limit for such analysis.

### Software

Image processing was carried out using Matlab (R_2023a). Data processing and analysis were performed in Python (CPython 3.10.11, IPython 8.12.0) with NumPy version 1.24.3, Pandas version 1.5.3 and iqplot 0.3.3 using Jupyter notebook (Jupyerlab version 3.6.3). Averages in Fig 4 were calculated from means of three different experiments and the error bar denotes the standard deviation between the experiments using .mean and .std methods of pandas dataframe respectively. Data was plotted with Bokeh version 3.1.1 and the figures were assembled with BioRender.com and Adobe Illustrator.

### Cryo-ET

#### Cryo-ET Sample Preparation

200 μL of 10 nm gold fiducial beads (Aurion) were centrifuged with a benchtop centrifuge for 20 minutes at 15,000 RPM and buffer exchanged with HEPES-NaOH buffer, pH 7.5. This procedure was repeated twice, and the beads were resuspended in a final volume of 100μL of HEPES-NaOH buffer, pH 7.5. Afterwards, 2μL of gold fiducial beads were added to 4μL of each sample. The droplet samples for Cryo-ET observation were prepared as previously, but with the following differences: DNA was incubated with HEPES and gold beads for 7 min, followed by addition of unlabeled P700. Droplets were induced by addition of Mg^2+^ and spotted on the grids after one minute of droplet induction. Water was used as a control for the no DNA case. Final concentrations: 1 mg/mL P700 (unlabeled), ∼50mM HEPES-NaOH, Mg^2+^: 100mM, DNA: [0-100 μg/mL].

Quantifoil R2/1 copper 200-mesh grids were glow-discharged with a Pelco easiGlow using the following parameters: set-15 mA, glow-25 seconds, and hold-10 seconds. 4μL of the samples containing the fiducial beads were deposited onto the grid and plunge-frozen into a propane/ethane mixture using a Vitrobot (Thermo Fisher Scientific) with the following parameters: 2.5 seconds blot time, 0 seconds wait time, 0.5 seconds drain time, 0 blot force, and 1 blot total.

#### Data Collection and Reconstruction

Cryo-ET samples were imaged using a 300 keV transmission electron microscope, Titan Krios (Thermo Fisher Scientific), equipped with a Gatan K3 direct electron detector and an energy filter (slit width of 20 eV was used). The data collection package SerialEM^112^ was used to run PACEtomo^113^ for tilt series acquisition. 35 image stacks were collected from −51° to +51° for each tilt series with an increment of 3°, a target defocus of −6 μm, a pixel size of 1.67 Å/pixel, and a total dose of approximately 100 e−/Å^2^. Each stack contained 10 frames, which were aligned using Motioncorr2^114^ and then assembled into the drift-corrected stack using IMOD. The drift-corrected stacks were aligned using fiducial markers and reconstructed by IMOD^115^.

#### Subtomogram Averaging

The subtomogram averaging package I3^116^ (version 0.9.9.3) was used to average the condensate edges. For each tomogram, the coordinate of the center of the condensate and multiple coordinates of the condensate edges were manually selected: polyP (2231 particles), polyP + pUC19 (1847 particles), polyP + pUC19 (10x) (1791 particles), polyP + 15 kb (1726 particles). An in-house script was used to calculate the Euler angles to orient particles in a consistent orientation. Subtomogram averaging was performed using bin4 particles reconstructed in the Weighted Back-Projection (WBP) method. The “graph” function in IMOD was used to generate density profiles.

### 3D Segmentation and Visualization of Cryo-ET Data

The representative tomograms shown in figure panels have been denoised by IsoNet^117^. 3D segmentations were generated using Dragonfly (2022.2) Deep Learning software (Object Research Systems)^118^. A 2D U-Net model was trained on an individual tomogram using hand-segmented frames of the corresponding tomogram. The model was then applied to the tomogram to generate a full 3D segmentation of the tomogram and then manually corrected. This process was repeated for each tomogram shown in Fig 3. The model was trained iteratively to distinguish the polyP interior, the dense edge of the condensate, the extruding DNA, and the background. Due to an inability to fully distinguish the dense edge and tightly wound DNA, the dense edge feature was depicted in yellow as shown in Fig 3f-h. Videos and 3D rendering images shown in figure panels were generated using UCSF ChimeraX (1.6.1)^119^.

## ACKNOWLEDGMENTS

We gratefully acknowledge support from the US NIH (NIGMS Grant R35 GM130375 to A.A.D. and Grant DP2-GM-739-140918 to L.R.R), Scripps Research start-up funds (to D.P.), and a Postdoctoral Fellowship from the American Heart Association (Award #903967 to R.C.). D.A.G is supported by the Pew Scholars Program. We would like to thank Anthony Milin and Ya-Ting Chang for help with preliminary studies. We would also like to thank Megan Bergkessel, Keren Lasker and Samrat Mukhopadhyay for insightful feedback on this work, and The Scripps Research Institute Core Microscopy Facility for use of confocal microscopy instrumentation.

