## Supplementary Information for "Reentrant DNA shells tune polyphosphate condensate size"

**SI Movie 1.** PolyP-$Mg^{2+}$ condensates form spherical droplets that fuse over time. (1mg/mL PolyP, 10% PolyP-AF647, 100 mM $MgCl_2$, 50mM HEPES, pH 7.5; scale bar = 10μm)

**SI Movie 2.** PolyP-$Mg^{2+}$ condensates form vacuoles under non-equilibrium conditions that fuse with time (1mg/mL PolyP, 300 mM $MgCl_2$, 50mM HEPES, pH 7.5; scale bar = 10μm)

**SI Movie 3.** PolyP-$Mg^{2+}$-DNA condensates form spherical PolyP-$Mg^{2+}$ condensates (PolyP-AF647, blue) surrounded by a DNA shell (1μM YOYO1, yellow) (1mg/mL PolyP, 10μg/mL pUC19, 100 mM $MgCl_2$, 50mM HEPES, pH 7.5; scale bar = 10μm)

**SI Movie 4.** Visualization of PolyP-$Mg^{2+}$ condensate on cryo-EM grid. Top-down and side views of the 3D rendering shown in Figure 3h. PolyP is shown in dark red, the PolyP dense edge is shown in light red, the carbon/PolyP interface is indicated by a green bracket, and DNAs are shown in cyan.

**SI Movie 5.** Visualization of PolyP-$Mg^{2+}$-pUC19 condensate on cryo-EM grid. Top-down and side views of the 3D rendering shown in Figure 3i. PolyP is shown in dark red,the dense edge+DNA is shown in yellow, the carbon/PolyP interface is indicated by a green bracket, and DNAs are shown in cyan.

**SI Movie 6.** Visualization of PolyP-$Mg^{2+}$-pUC19(10X) condensate on cryo-EM grid. Top-down and side views of the 3D rendering shown in Figure 3j. PolyP is shown in dark red, the dense edge+DNA is shown yellow, the carbon/PolyP interface is indicated by a green bracket, and DNAs are shown in cyan.

**SI Movie 7.** Visualization of PolyP-$Mg^{2+}$-15kb condensate on cryo-EM grid. Top-down and side views of the 3D rendering shown in Figure 3k. PolyP is shown in dark red, the dense edge+DNA is shown in yellow, the carbon/PolyP interface is indicated by a green bracket, and DNAs are shown in cyan.

**SI Movie 8.** Addition of 15kb plasmids, results in smaller DNA-shelled condensates (PolyP-AF647: blue, 1μM YOYO1: yellow) (1mg/mL PolyP, 10μg/mL pUC19, 100 mM $MgCl_2$, 50mM HEPES, pH 7.5; scale bar = 10μm).

**SI Movie 9.** When T4 DNA was added, grape-like clusters of condensates emerged. These clusters moved together as a unit, but did not fuse on the timescale observed.

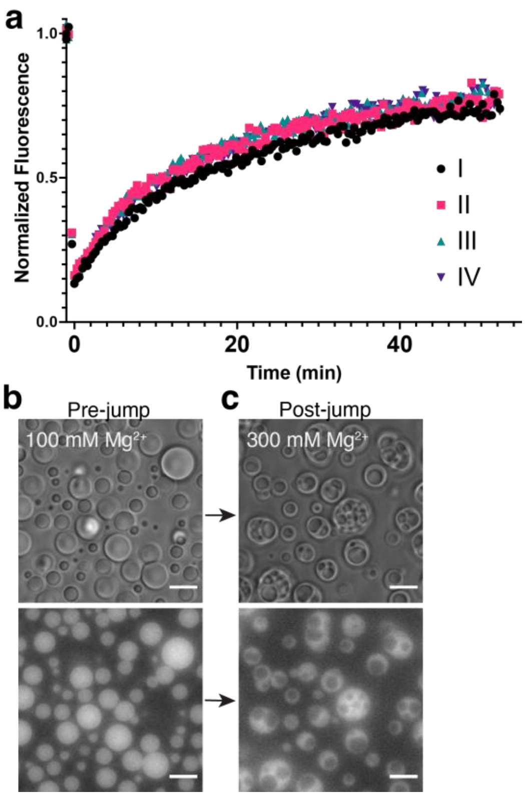


**Fig S1.** a) Overlay of individual FRAP experiments for PolyP-$Mg^{2+}$ condensates. Condensates were allowed to grow for 35-45 minutes after which a small circular region in the center of the condensate was bleached. Scans were taken every 20s for 52 minutes with autofocus z corrections being applied every 15 images (~5 min). New samples were prepared for each experiment. Each run takes place between 35-100min after droplet formation. The time zero on the graph is the scan immediately following the bleach. b) PolyP-$Mg^{2+}$ condensates form vacuoles upon addition of $Mg^{2+}$ to preformed droplets. PolyP-$Mg^{2+}$ condensates were formed (1 mg/mL PolyP, 100mM $MgCl_2$, 50 mM HEPES pH 7.5) and allowed to fuse and grow for 10 min. c) $Mg^{2+}$ concentration was subsequently brought to 300mM. Within a minute of $MgCl_2$ addition, vacuoles were observed. These vacuoles fused (Mov. S1) and were transient.

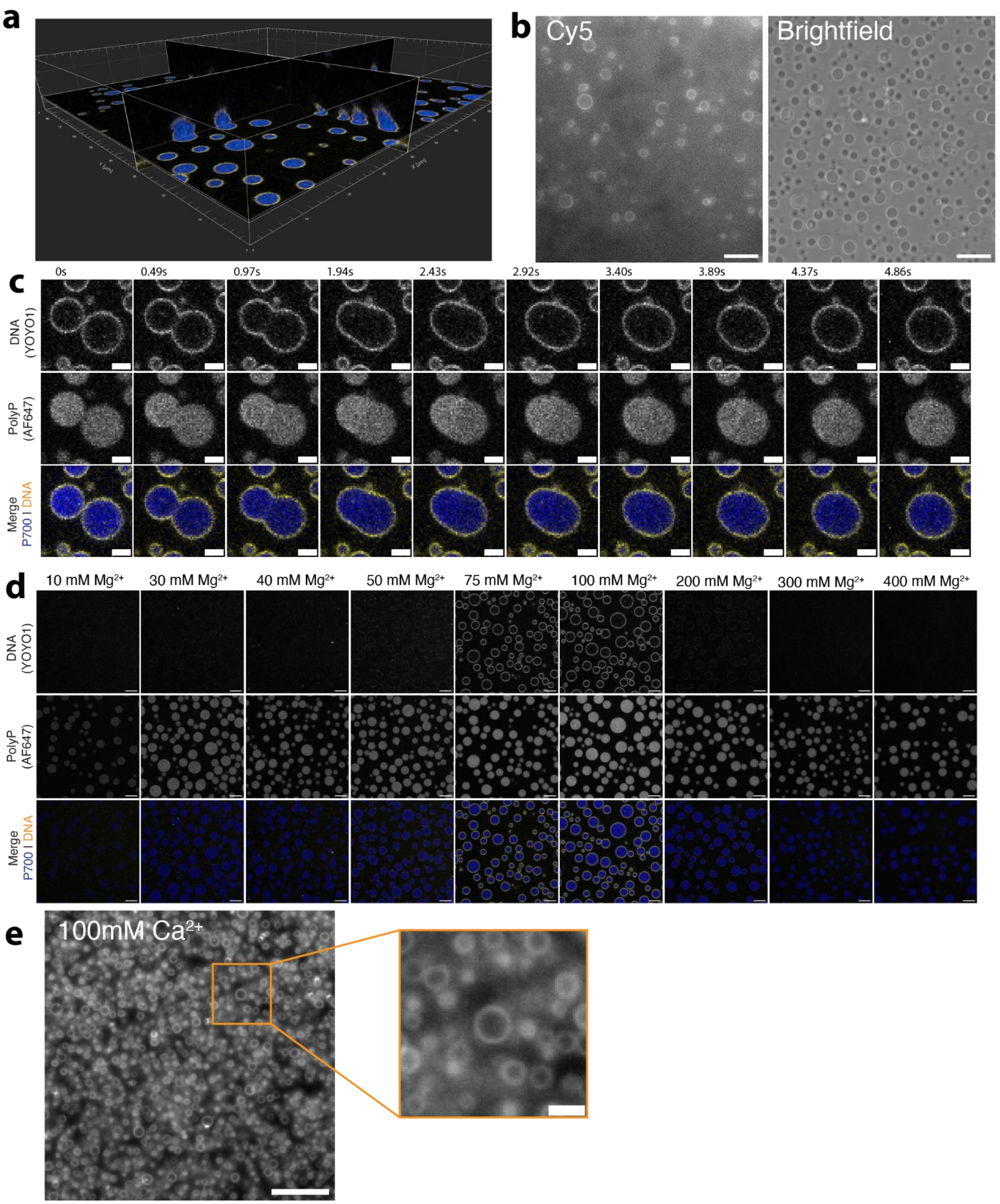


**Fig. S2.** a) Imaris-rendered ortho slices of PolyP-$Mg^{2+}$-pUC19 shelled condensates (1mg/mL PolyP (10% P700-AF647, blue), 100mM $Mg^{2+}$, 10ng/uL pUC19 (1µM YOYO1, yellow)). b) DNA shells visualized using 5' end-labeled Cy5 ([pUC19] = 5 µg/mL, scale bar = 10µm). c) Frame-by-frame visualization of P700-$Mg^{2+}$-pUC19 condensate fusion (scale bar = 2µm). d) DNA shells show reentrant behavior, appearing between 50mM and 200mM $Mg^{2+}$ (scale bar = 5µm). e) P700 condensates also form in the presence of $Ca^{2+}$ (1mg/mL PolyP, 100 mM $CaCl_2$, scale bar (main) = 10µm, scale bar (inset) = 2µm ).

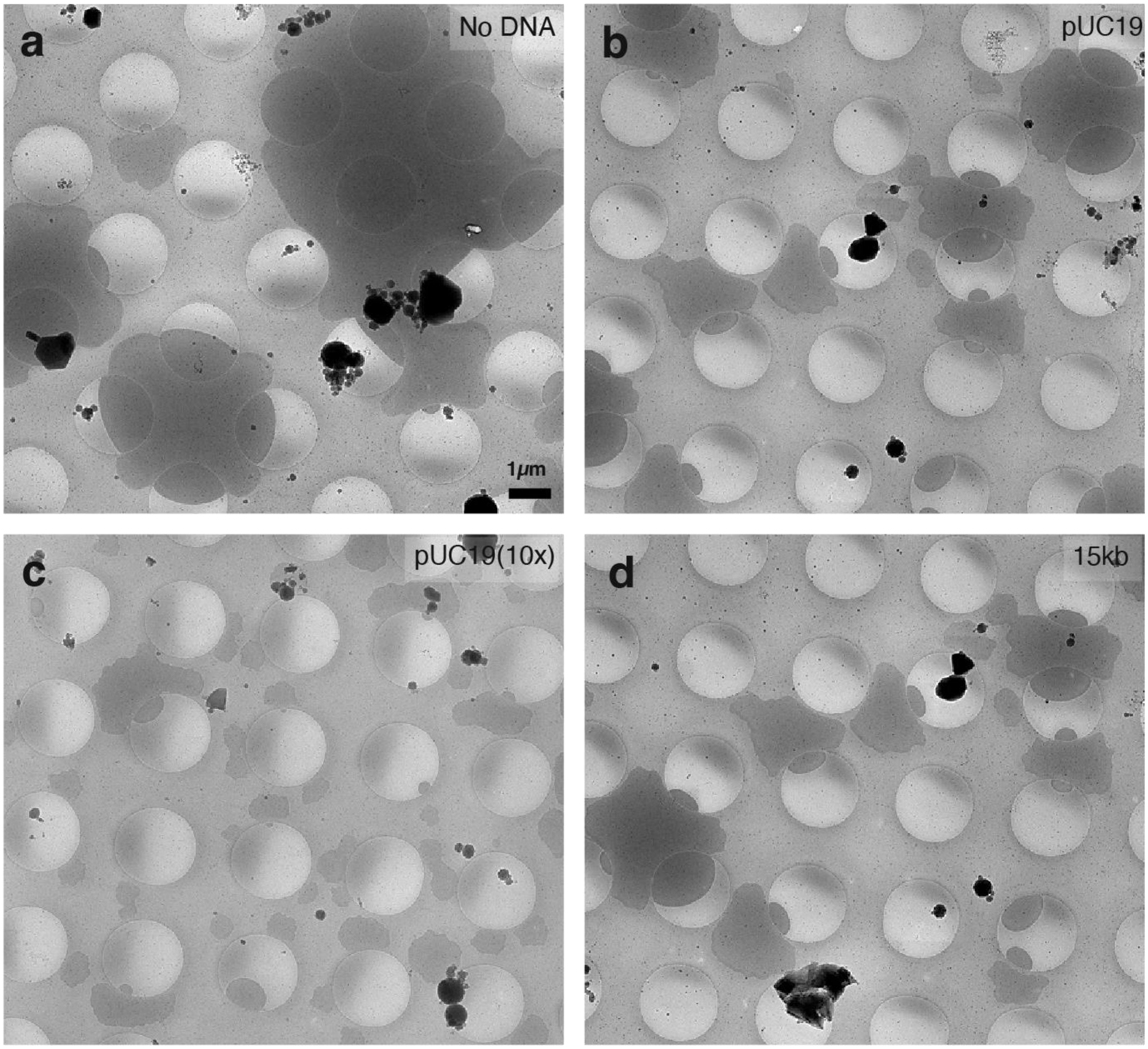


**Fig. S3.**(a-d) Low-magnification cryo-electron microscopy (cryo-EM) images of PolyP-$Mg^{2+}$ condensates with various types of DNA on cryo-EM grids: (a) No DNA (b) pUC19, (c) pUC19 (10x), (d) 15kb.

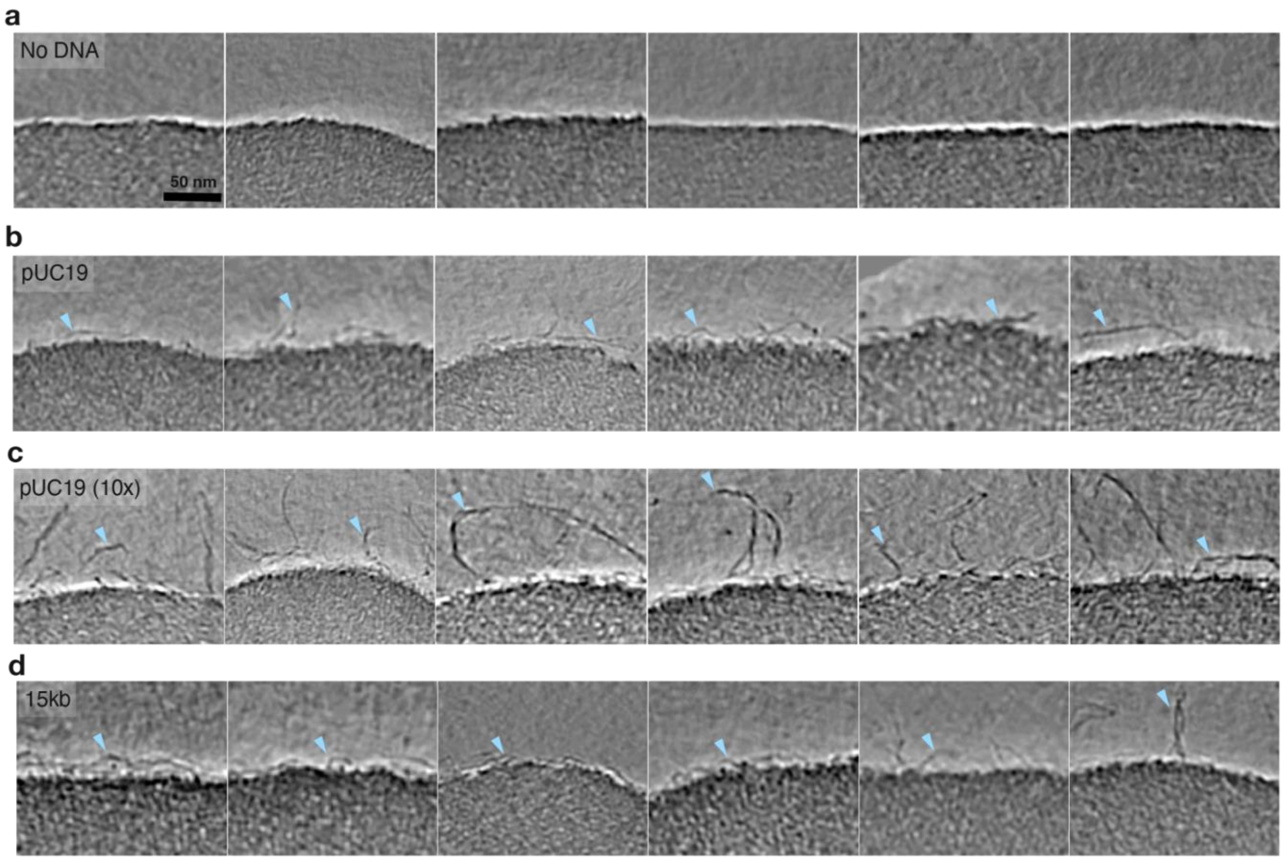


**Fig S4.** Gallery of snapshots showing the surface of PolyP condensates incubated with different types of DNA. Cyan arrows highlight DNA that sticks out of the surface

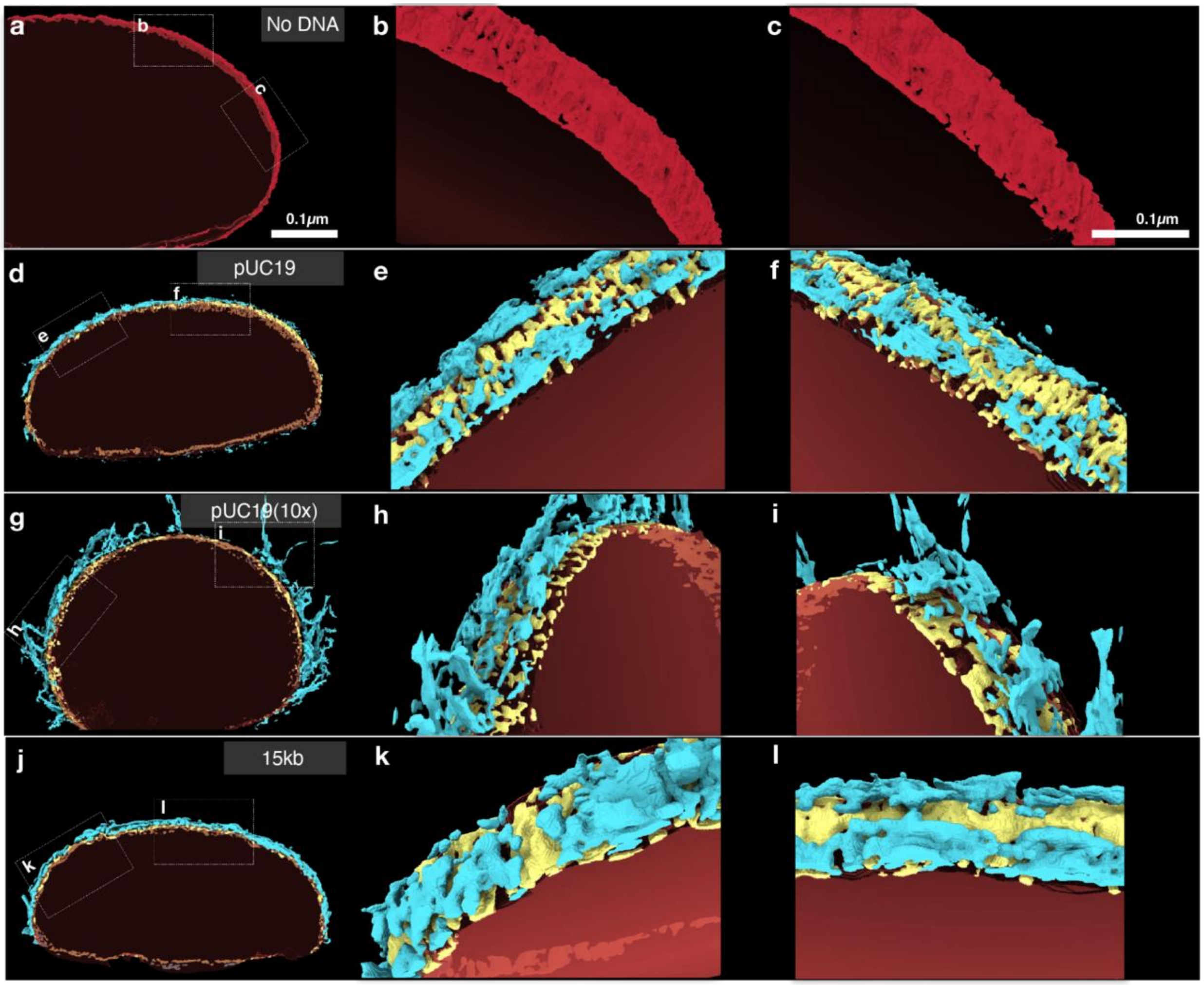


**Fig. S5.** Surface texture of PolyP-$Mg^{2+}$ condensates interacting with different types of DNAs: (a-l) 3-dimensional renderings of tomograms shown in Figure 3 in both top-down views (a, d, g, j) and surface views (b-c, e-f, h-i, k-l). PolyP condensate edges are shown in red, DNA filaments are in cyan, and the ambiguous polyP-$Mg^{2+}$ dense edge and DNA surface is in yellow.

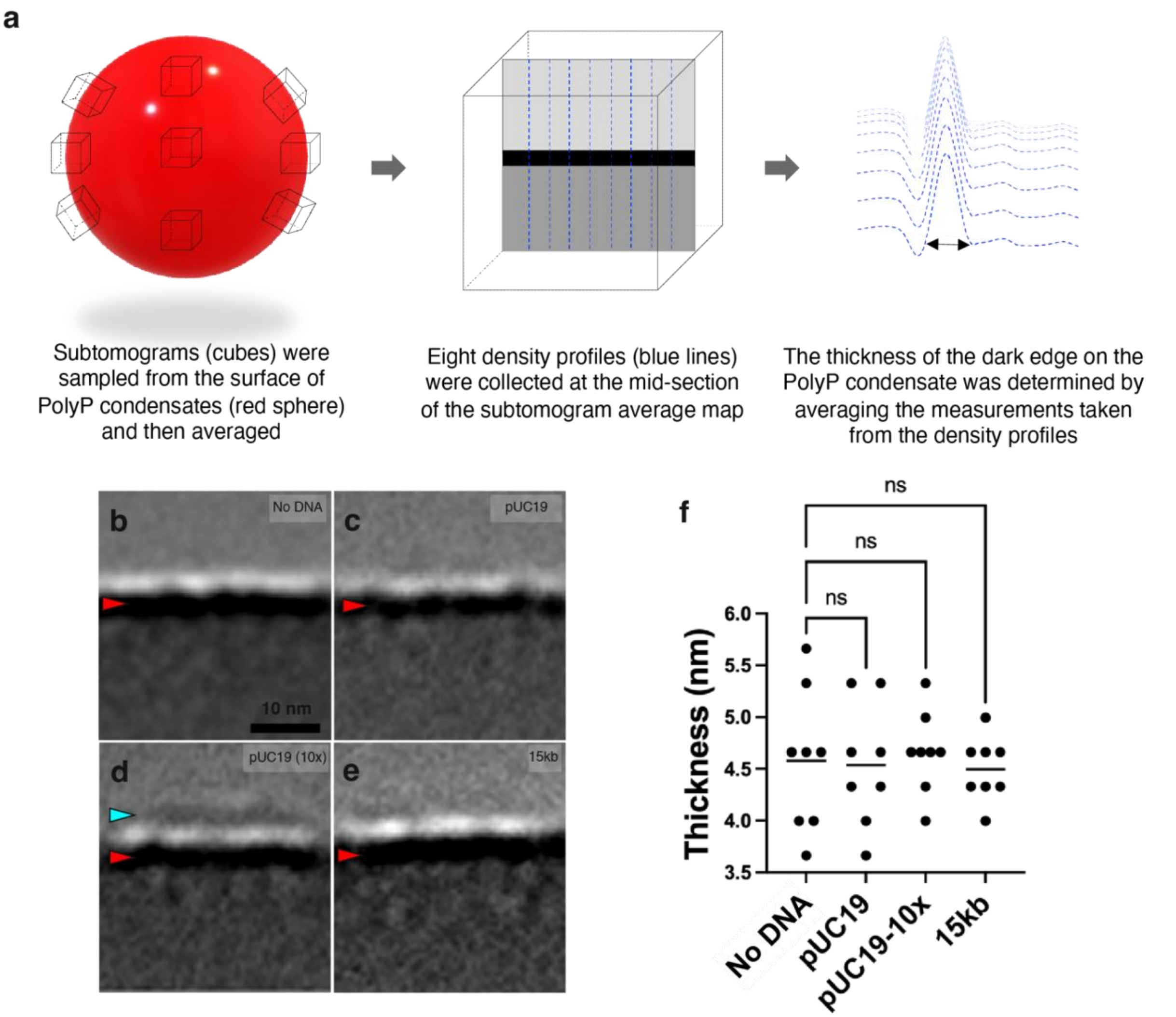


**Fig. S6.** (a) A cartoon model depicting the subtomogram sampling process and the thickness measurement. (b-e) Subtomogram averages of the surface of PolyP condensates incubated with different types of DNA. Red arrows indicate the dense edges of condensates, and the cyan arrow in panel e indicates extra density found only in this experimental condition. (f) Comparison of condensate edge thickness. The thickness of the dark edges was measured as described in Panel a. The measured thickness values for the different samples are as follows: (b) 4.6±0.7 nm, (c) 4.5±0.6 nm, (d) 4.7±0.4 nm, (e) 4.5±0.3 nm, Statistical significance was calculated using one-way ANOVA with the Tukey HSD multiple comparison test.

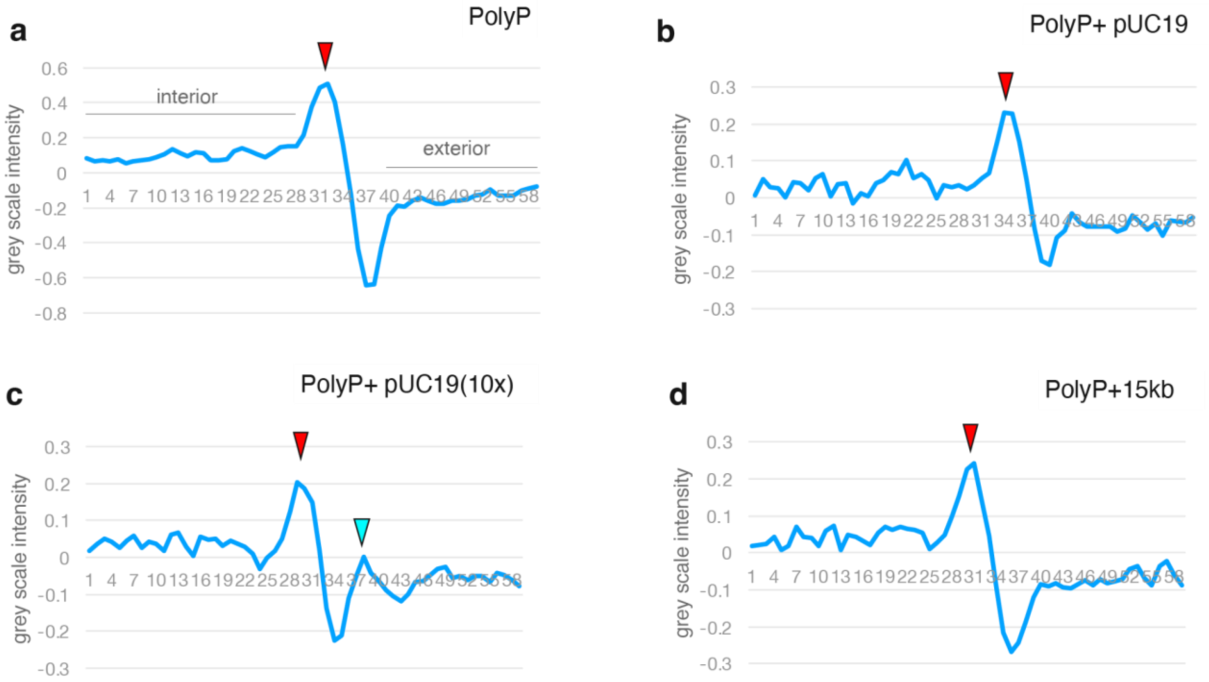


**Fig. S7.** (a-d) Representative density profiles. The x-y plane density profile is drawn perpendicular to the direction of the dense edge. The X-axis represents the vertical span of pixels (6.65Å/pixel), extending from the bottom to the top of the average map. The Y-axis represents grayscale values ranging from -1 (white) to +1 (black). Red arrows indicate the dense edges of condensates, and the cyan arrow in panel d indicates extra density found only in this experimental condition.

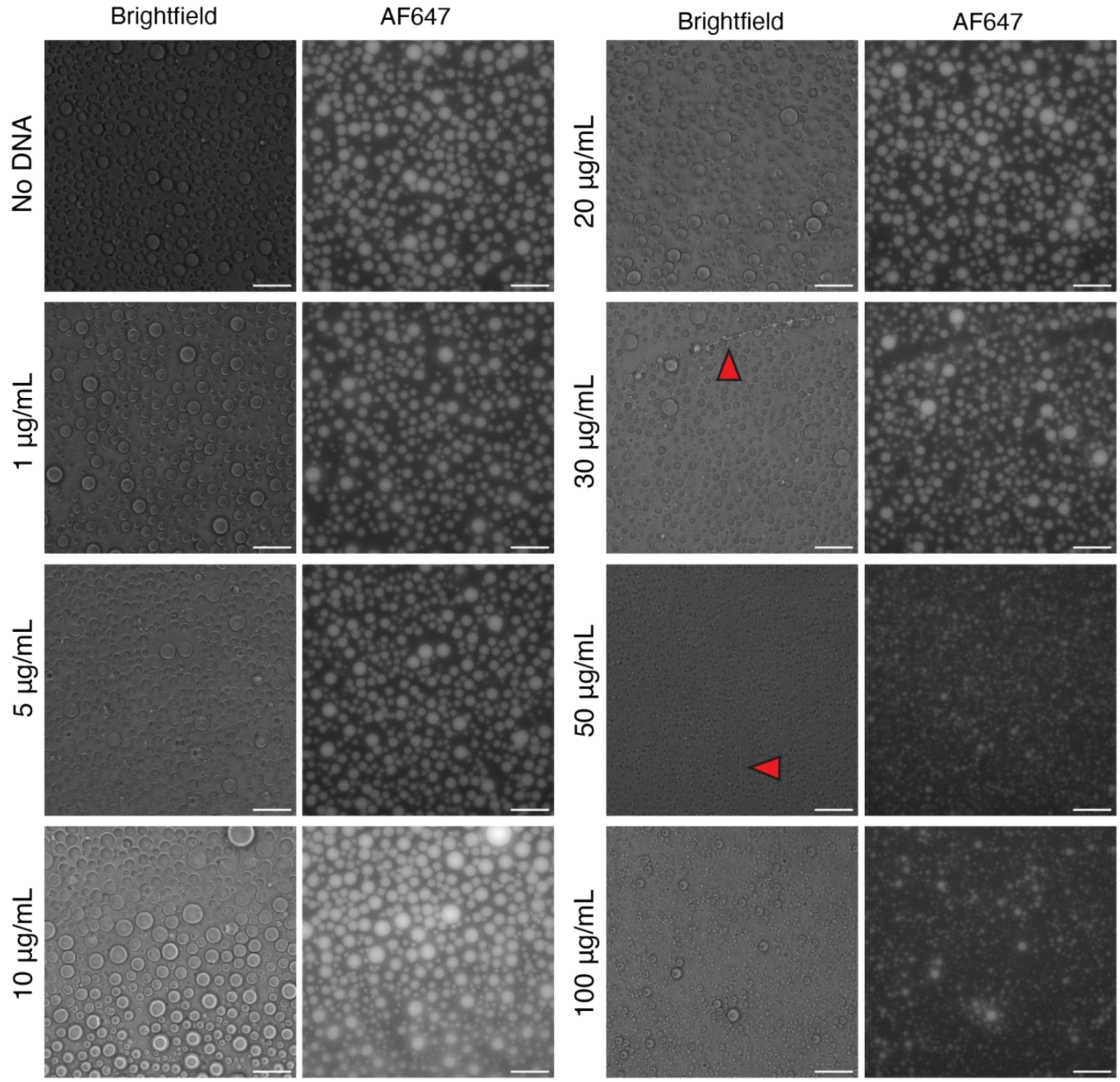


**Fig. S8.** Representative widefield images in brightfield and fluorescence detection channels with varied DNA concentrations used for droplet size analysis (1mg/mL PolyP, 100mM MgCl2, 50mM HEPES; scale bar = 20µm). Filaments were observed in some fields of view above 30µg/mL DNA. Those visualized in these fields of view are indicated by red arrows.

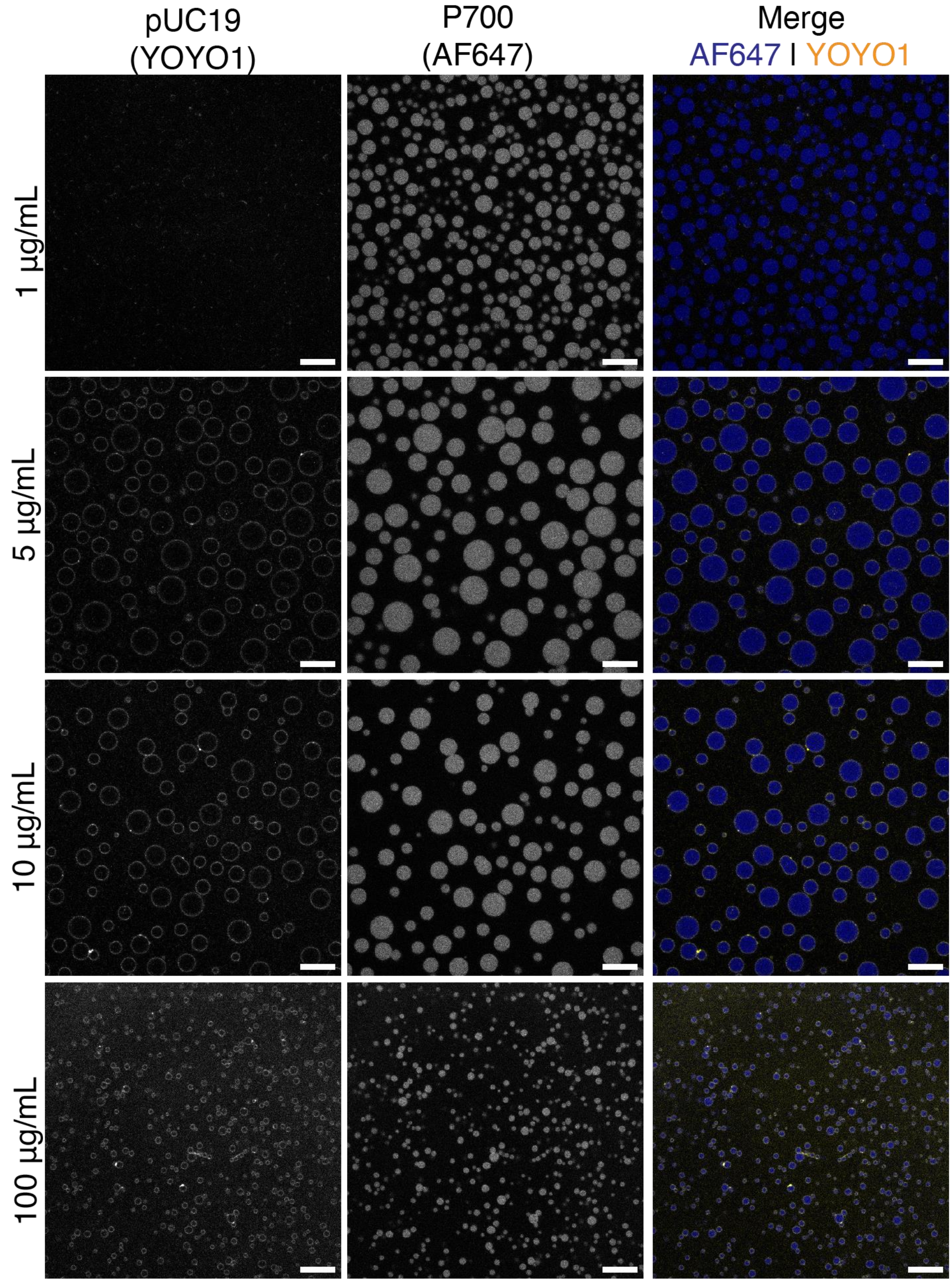


**Fig. S9.** Representative confocal images of varied pUC19 concentration (1mg/mL PolyP (10% P700-AF647), 100mM MgCl2, 50mM HEPES, 1μM YOYO1; scale bar = 10μm).

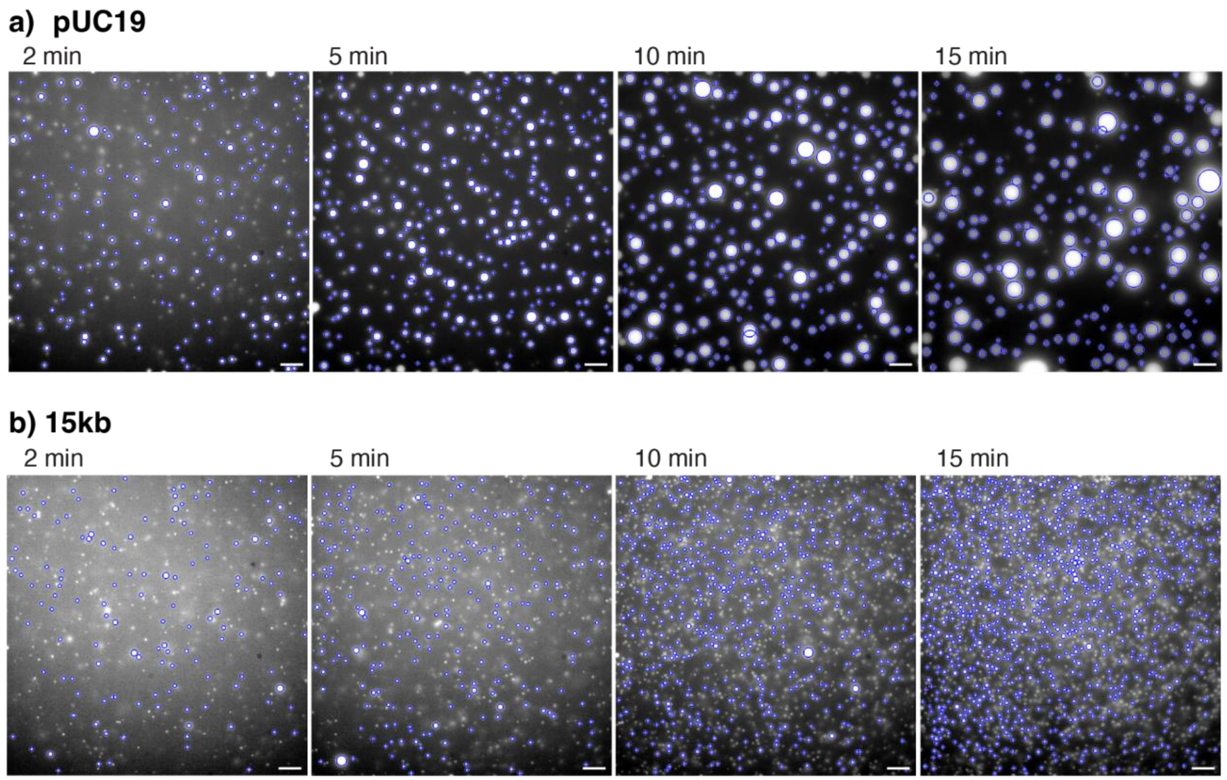


**Fig S10**. Representative overlays from segmentation step using the widefield 640 channel images for a) pUC19 and b) 15kb condensates. Images shown here were selected from the first of four fields of view from a single experiment at 2, 5, 10, and 15 min time points.

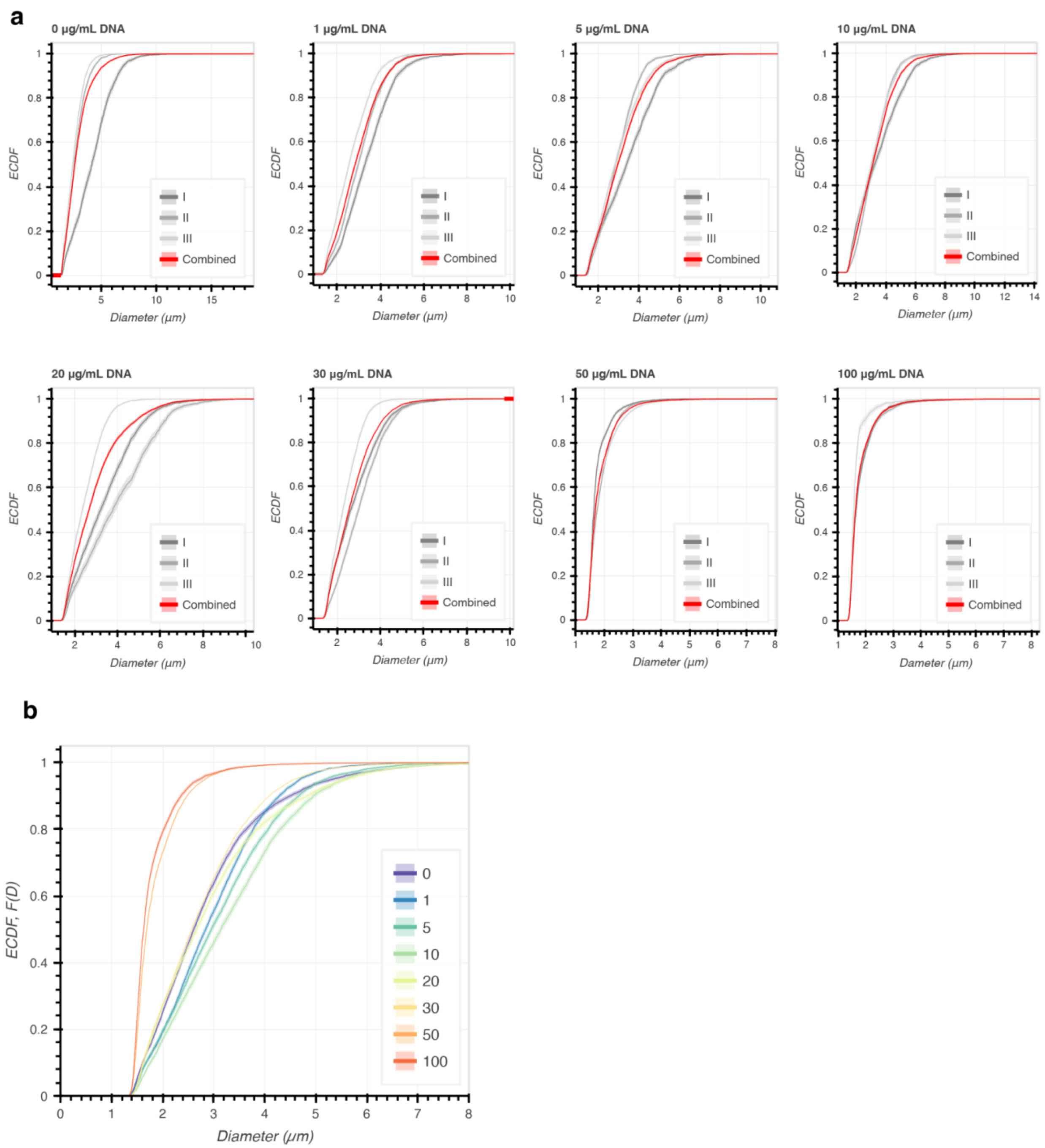


**Fig S11.** a) ECDF curves showing cumulative distributions for varied DNA concentrations at 10 minutes for individual experiments (grey) and the combined distribution (red). Shaded regions behind the solid line represent 95% confidence intervals calculated through Iqplot's bootstrapping methods. b) The combined distribution ECDFs shown in red in panel a are overlaid together for reference for the various DNA concentrations.

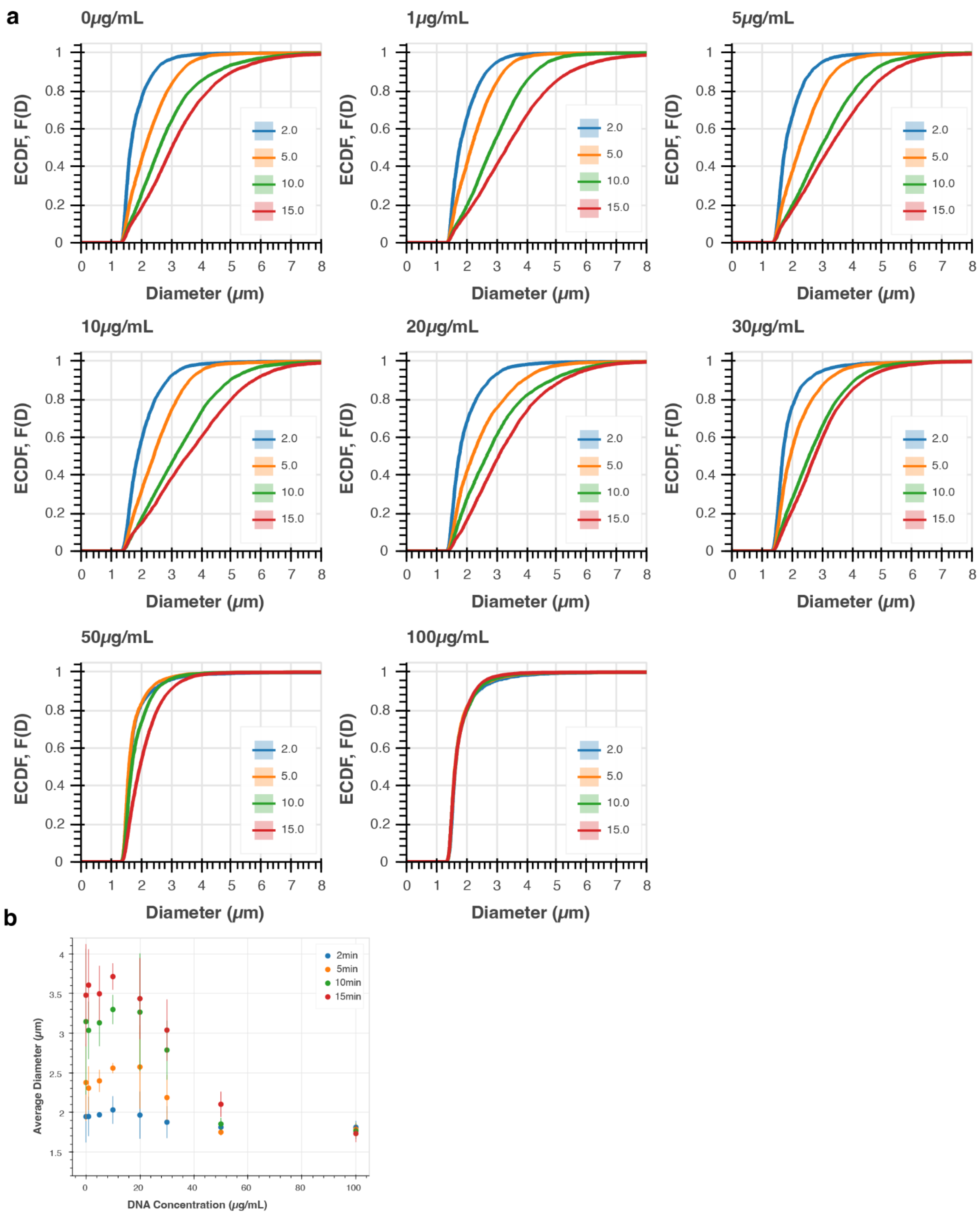


**Fig S12.** a) ECDF curves representing the cumulative distributions for varied DNA concentrations at 2, 5, 10, and 15 minute timepoints. b) Compiled average of average diameters for various DNA concentrations at 2, 5, 10, and 15 minute timepoints across three experiments. Error bars represent the standard deviation from the mean of mean diameter for the three replicates.

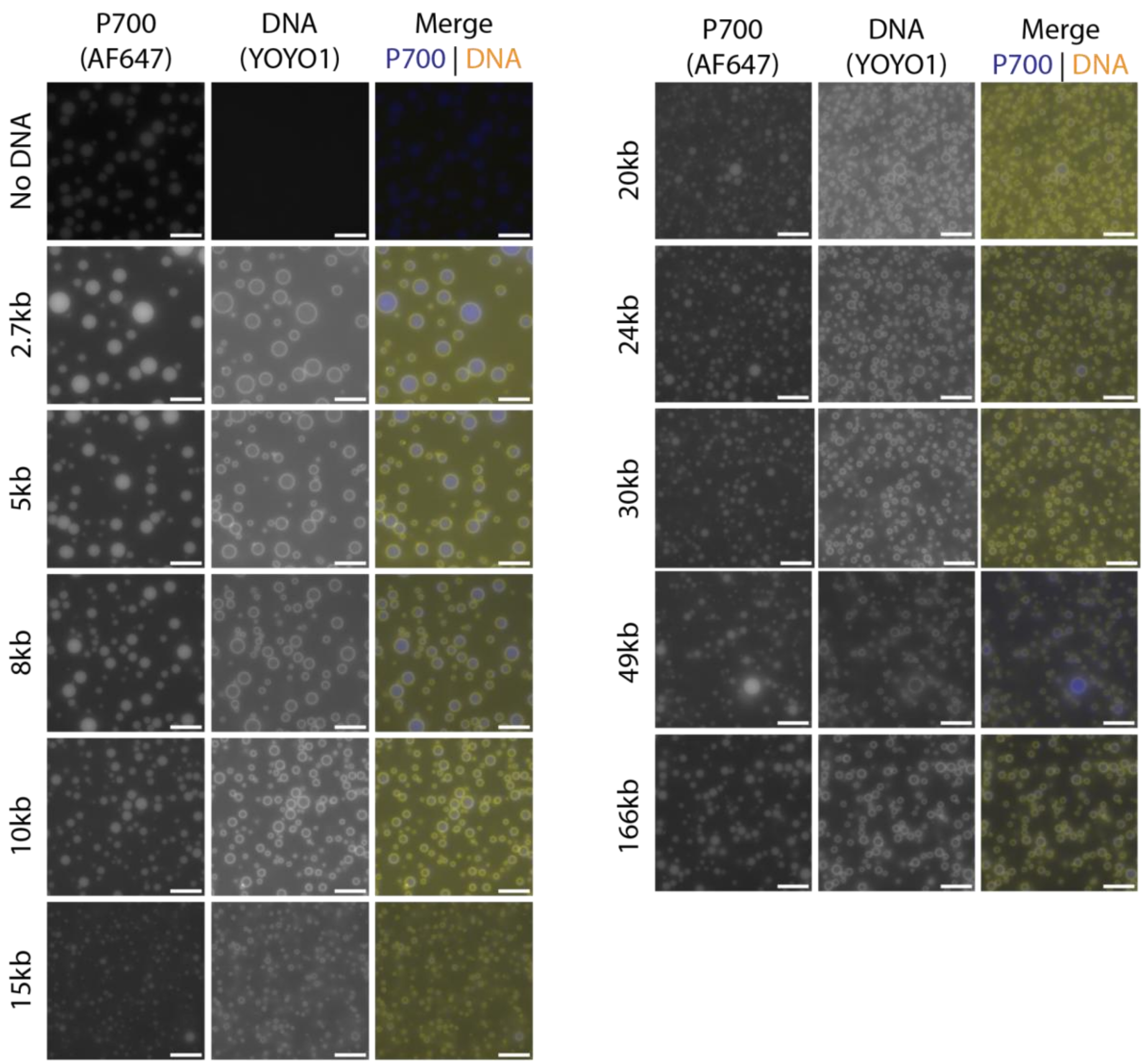


**Fig S13.** Representative widefield images with varied DNA lengths (1 mg/mL P700 (10% P700-AF647), 100mM $MgCl_2$, 50mM HEPES,1 μM YOYO1, 10ng/μL DNA; scale bar = 5μm). Individual channels are shown in gray, while the merge uses blue for PolyP (P700-AF647) and yellow for DNA (YOYO1).

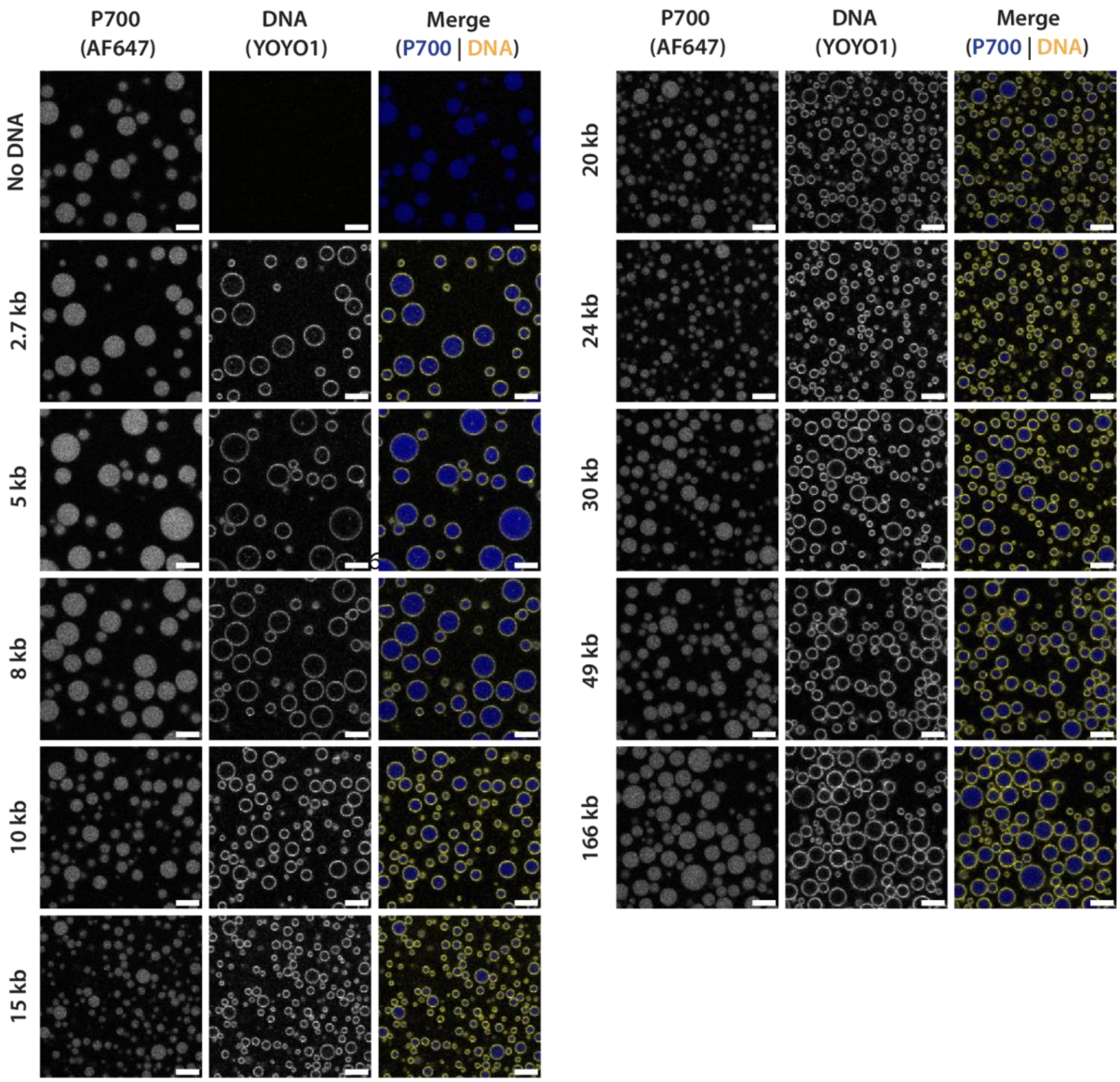


**Fig S14.** Representative confocal images with varied DNA lengths (1 mg/mL P700 (10% P700-AF647), 100mM $MgCl_2$, 50mM HEPES,1 μM YOYO1, 10ng/μL DNA;  scale bar = 5μm)

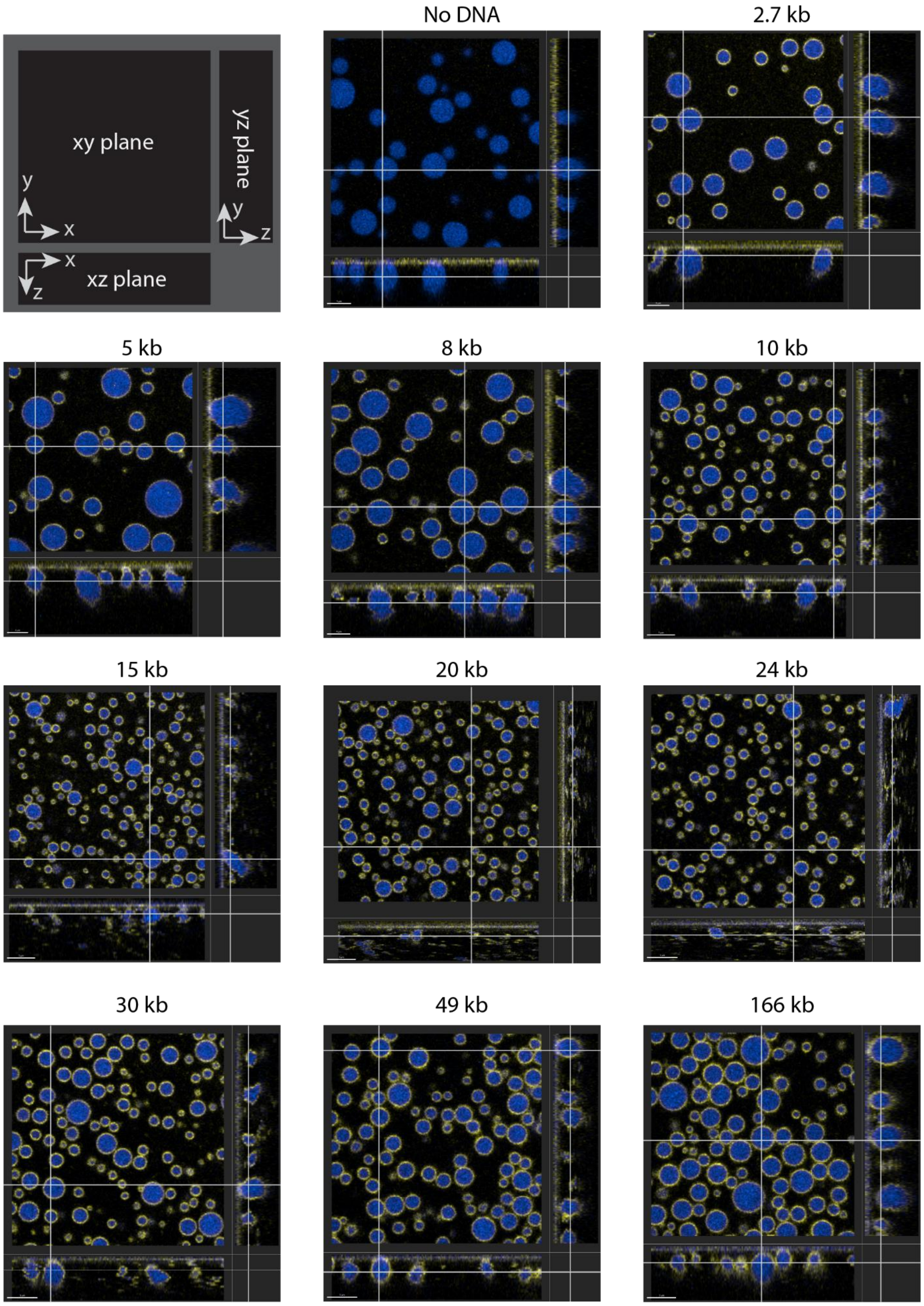


**Fig S15.** Imaris Orthoview renderings of PolyP-$Mg^{2+}$-DNA confocal z-stacks for varied DNA length experiments (1 mg/mL P700 (10% P700-AF647), 100mM MgCl2, 50mM HEPES,1 µM YOYO1, 10ng/µL DNA;  P700: blue, DNA: yellow; scale bar = 5µm). Z-stacks were collected at 10 minutes after condensate formation with 0.37µm between frames, except for 20 & 24kb which were collected at  0.15µm and 30kb & 48kb which were at 0.30µm.

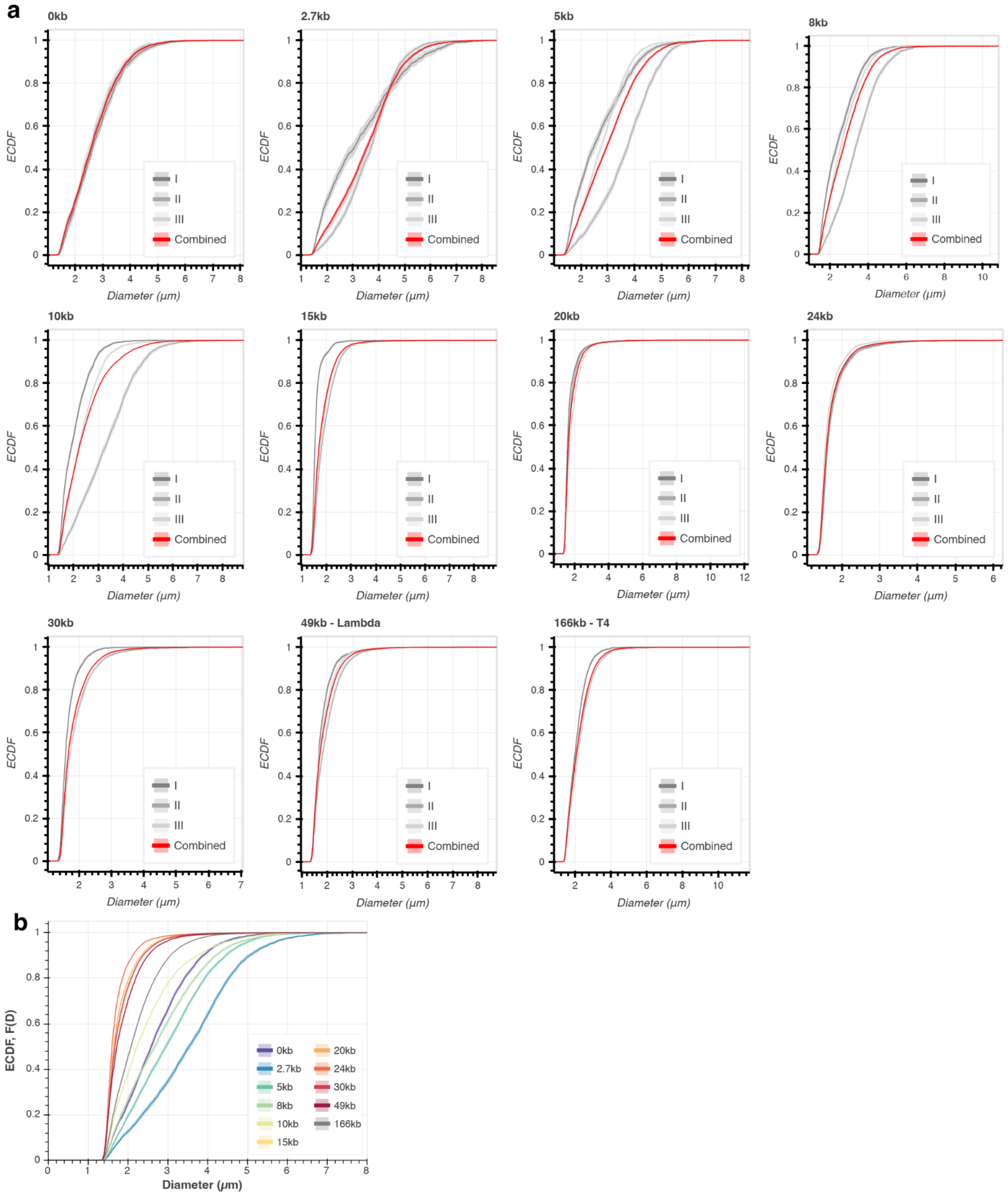


**Fig S16.** a) ECDF curves showing cumulative distributions for varied DNA lengths at 10 minutes for individual experiments (grey) and the combined distribution (red). Shaded regions behind the solid line represent 95% confidence intervals calculated through Iqplot's bootstrapping methods. b) The combined distribution ECDFs shown in red in panel a are overlaid together for reference for the various DNA lengths.

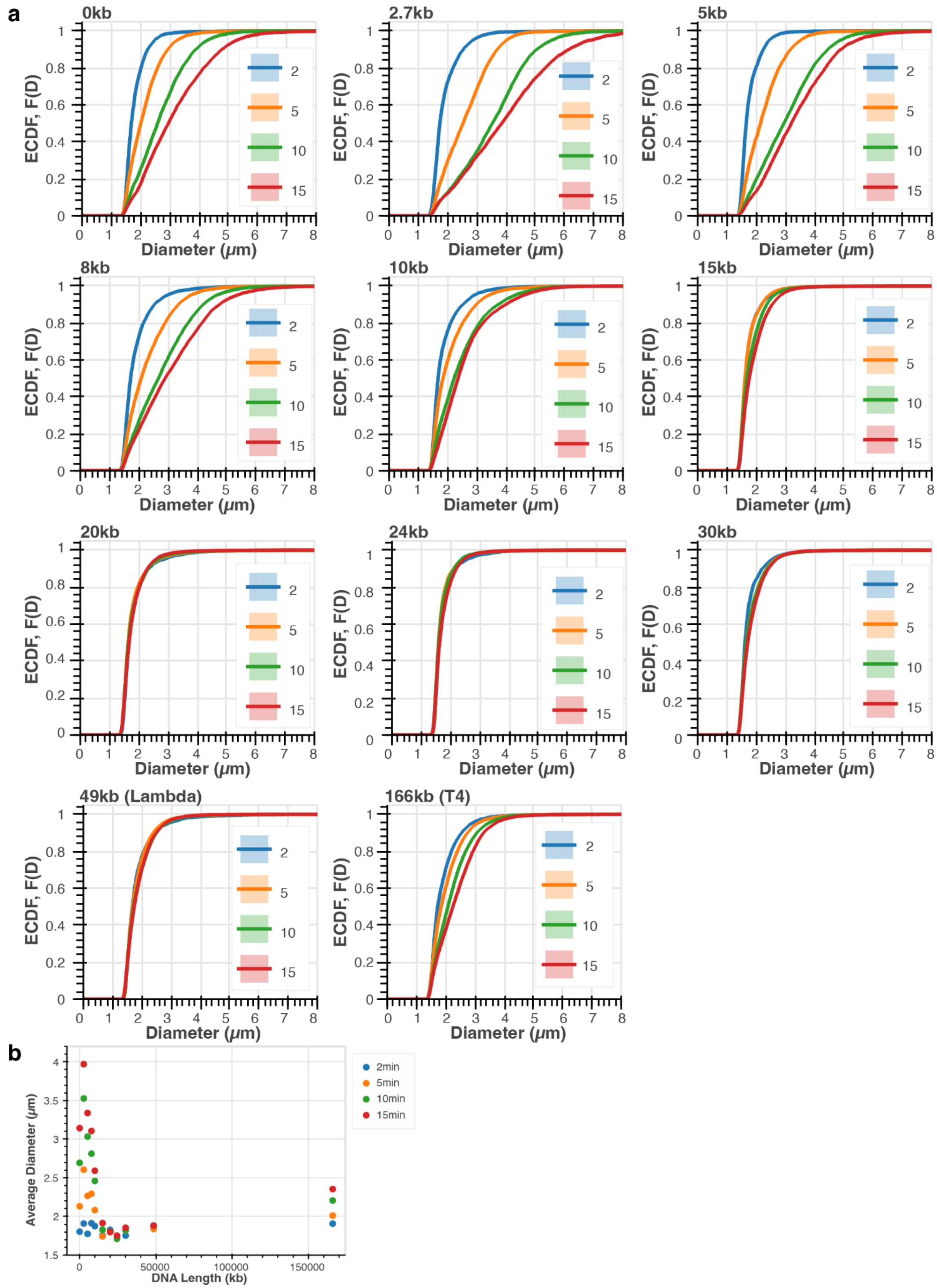


**Fig S17**. a) ECDF curves representing the distributions with varied DNA length from compiled experiments. b) Compiled average diameters for various DNA lengths at 2, 5, 10, and 15 minute timepoints.
